# A Comprehensive Assessment of Demographic, Environmental and Host Genetic Associations with Gut Microbiome Diversity in Healthy Individuals

**DOI:** 10.1101/557124

**Authors:** Petar Scepanovic, Flavia Hodel, Stanislas Mondot, Valentin Partula, Allyson Byrd, Christian Hammer, Cécile Alanio, Jacob Bergstedt, Etienne Patin, Mathilde Touvier, Olivier Lantz, Matthew L. Albert, Darragh Duffy, Lluis Quintana-Murci, Jacques Fellay, The *Milieu Intérieur* Consortium

## Abstract

**Background:** The gut microbiome is an important determinant of human health. Its composition has been shown to be influenced by multiple environmental factors and likely by host genetic variation. In the framework of the *Milieu Intérieur* Consortium, a total of 1,000 healthy individuals of western European ancestry, with a 1:1 sex ratio and evenly stratified across five decades of life (age 20 – 69), were recruited. We generated 16S ribosomal RNA profiles from stool samples for 858 participants. We investigated genetic and non-genetic factors that contribute to individual differences in fecal microbiome composition.

**Results:** Among 110 demographic, clinical and environmental factors, 11 were identified as significantly correlated with α-diversity, ß-diversity or abundance of specific microbial communities in multivariable models. Age and blood alanine aminotransferase levels showed the strongest associations with microbiome diversity. In total, all non-genetic factors explained 16.4% of the variance. We then searched for associations between >5 million single nucleotide polymorphisms and the same indicators of fecal microbiome diversity, including the significant non-genetic factors as covariates. No genome-wide significant associations were identified after correction for multiple testing. A small fraction of previously reported associations between human genetic variants and specific taxa could be replicated in our cohort, while no replication was observed for any of the diversity metrics.

**Conclusion:** In a well-characterized cohort of healthy individuals, we identified several non-genetic variables associated with fecal microbiome diversity. In contrast, host genetics only had a negligible influence. Demographic and environmental factors are thus the main contributors to fecal microbiome composition in healthy individuals.

## BACKGROUND

A wide diversity of microbial species colonizes the human body, providing considerable benefits to the host through a range of different functions [1]. Notably, these microbes generate metabolites that can act as energy sources for cell metabolism, promote the development and the functionality of the immune system, and prevent colonization by pathogenic microorganisms [2].

The human intestine harbors a particularly diverse microbial ecosystem. Multiple 16S ribosomal RNA (rRNA) gene sequencing and metagenomic studies established that each individual gut microbiome harbors a unique combination of microbial life [3, 4]. An estimated 150 to 400 bacterial species reside in each person’s gut [5].

Typically, the human gut microbiome is dominated by five bacterial phyla: *Firmicutes, Bacteroidetes, Proteobacteria, Actinobacteria* and *Verrucomicrobia* [6, 7]. These contain almost all of the bacterial species found in the human gastrointestinal tract, which can also be classified in higher-level taxonomic groups such as genera, families, orders and classes [8]. The relative proportions of microbial species vary extensively between individuals [9] and has been shown to be age-dependent [10]. The microbiome composition evolves rapidly during the first three years of life, followed by a more gradual maturation [11], then is predicted to remain relatively stable throughout adult life [12].

A variety of environmental and clinical factors including diet, lifestyle, diseases and medications can induce substantial shifts in the microbiome composition [13, 14]. Multiple studies have shown that diet and medications are the main forces influencing gut microbial diversity [15, 16, 17, 18, 19, 20, 21, 22]. Yet, they only explain a small percentage of the microbiome variation observed in the human population. Host genetics has also been proposed as a contributor in determining the relative abundance of specific gut microbes [23, 24]. Several studies have searched for associations between human genetic variation and gut microbiome diversity [20, 21, 22, 25, 26, 27, 28], but only a few genetic loci have been replicated across these studies. As a consequence, most of the interindividual variability in gut microbiome composition remains unexplained.

In this study, we leveraged the in-depth phenotypic and genotypic information available for the *Milieu intérieur* (MI) cohort - a population-based study of 1,000 healthy individuals of western European ancestry, evenly stratified by sex (1:1) and age. We investigated the role of socio-demographic and environmental factors in inter-individual gut microbiome variation (Figure 1). In particular, we were able to assess the impact of family status, income, occupational status and educational level, smoking habits, sleeping habits, psychological problems, and nutritional behavior. We also evaluated the influence of basic physiological parameters (such as body mass index), family and personal medical history (including vaccination history) and multiple laboratory results (comprising mostly blood biochemical measurements). Finally, we investigated the potential impact of human genetic variation using a genome-wide association study (GWAS) framework, including as covariates the non-genetic factors that were found to be correlated with various measures of gut microbiome diversity.

**Figure 1.**
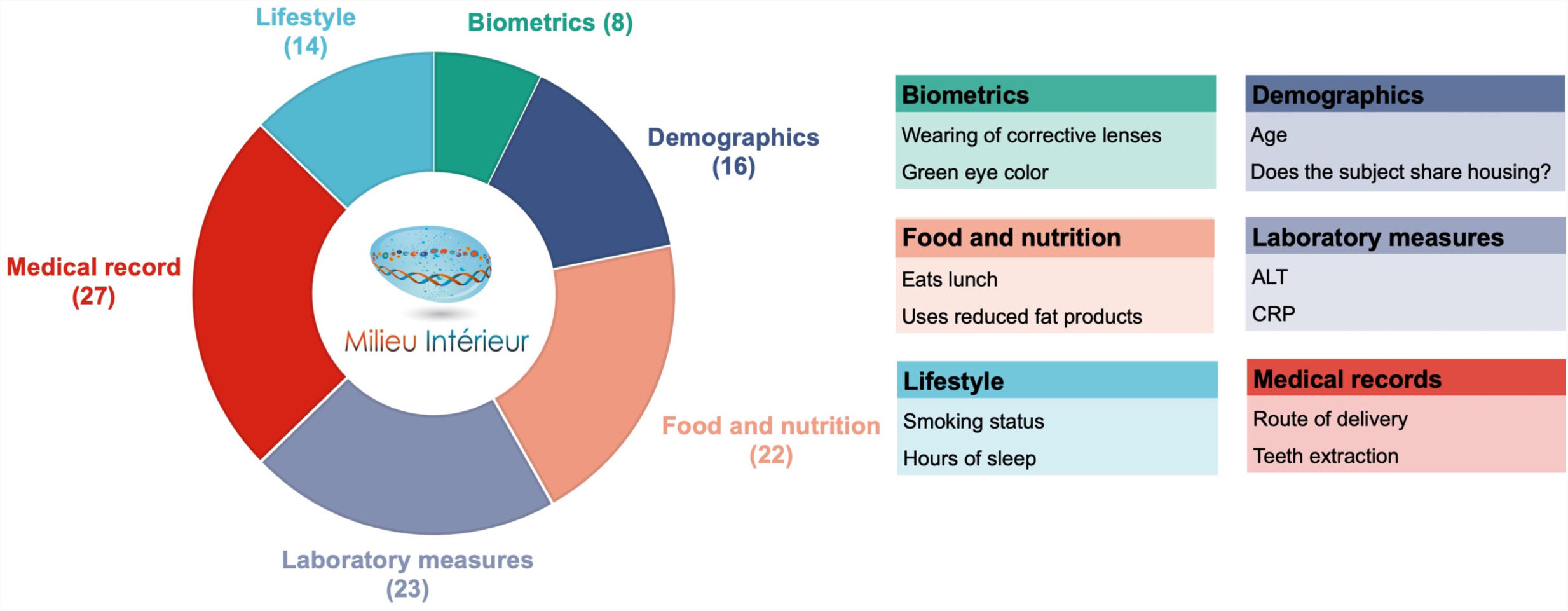
Non-genetic variables. Six categories of non-genetic variables investigated in this study. In the parenthesis, number of variables per each category and for each two representative examples. Full description of the variables is available in Additional File 2: Table S1.

## RESULTS

### Gut microbiome diversity in healthy donors

To characterize the bacterial diversity of the gut flora of the 1,000 healthy donors, we performed 16S rRNA gene sequencing on standardized collections of fecal samples. From this cohort, we obtained profiles for 858 individuals and we normalized the data for sequencing depth (see Methods). A total of 8,422 operational taxonomy units (OTUs) were detected, corresponding to 11 phyla, 24 classes, 43 orders, 103 families, 328 genera and 698 species. On average, we detected 193 species per individual (standard error 1.9, standard deviation 55.1), with a minimum of 58 and a maximum of 346 species. Inter-individual variability was already marked at the phylum level. Figure 2A presents the relative abundances of the 8 phyla observed in more than 10% of study participants. *Firmicutes* and *Proteobacteria* were detected in all individuals, and *Bacteroidetes* in all but one individual. *Firmicutes* was the dominant phylum in the vast majority of individuals (91.8%).

**Figure 2.**
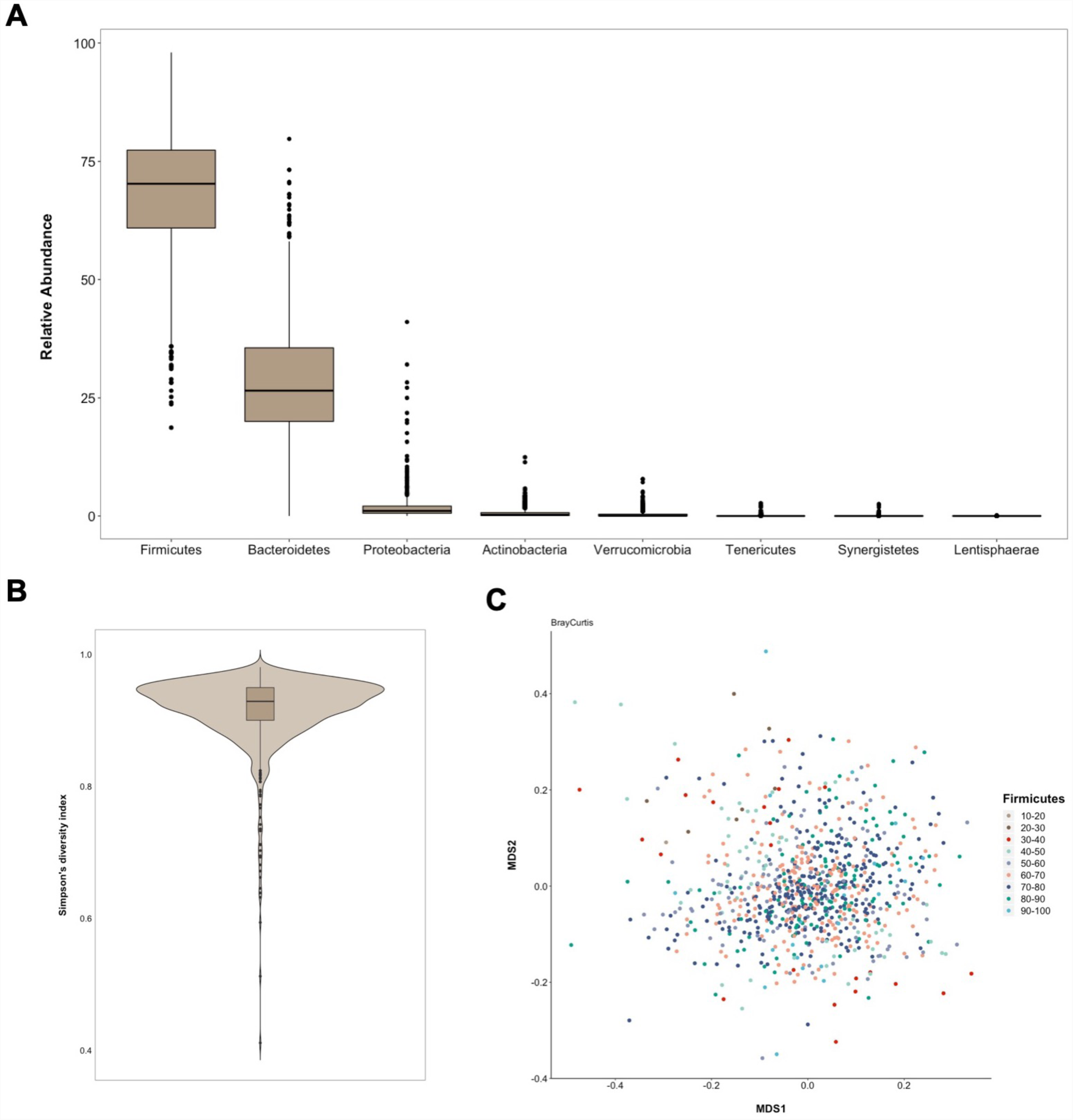
Gut microbiome diversity. (A) Box-plots of relative abundances of 8 phyla that were observed in more than 10% of the donors. Outliers are also represented. (B) Violin plot of Simpson’s diversity index values observed among MI study participants. (C) Multidimensional scaling plot of Bray-Curtis dissimilarity matrix with study participants colored according to relative abundance of *Firmicutes*.

Starting from the OTU counts, we calculated α and β microbiome diversity metrics (see Methods). As measures of α-diversity, which describes diversity within each sample, we used observed richness (number of distinct species present in the given sample), Chao1 richness estimate (estimate of the number of unobserved species), ACE (Abundance-based Coverage Estimator) and Simpson’s diversity index (probability that two randomly picked sequences belong to the same species). The histograms of their raw and transformed distributions are shown in Additional File 1: Figure S1A and S1B. We present here the results obtained using Simpson’s diversity index as a representative metric of α-diversity. The results for other indicated metrics are presented in the supplementary material. Figure 2B presents the distribution of Simpson’s diversity indexes depicting the continuous distribution and high diversity of the gut microbiome in the majority of study participants. The distributions of the other α-diversity metrics are shown in Additional File 1: Figure S1C.

As measures of β-diversity, which describes the difference in taxonomic composition between samples, we used compositional Jaccard (unweighted), as well as Bray-Curtis (weighed) and phylogenetic Unifrac (weighted) dissimilarity matrices. We present here the results obtained using Bray-Curtis dissimilarity matrix as a representative metric of β-diversity. The results for other indexes are presented in the supplementary material. Figure 2C presents the multidimensional scaling (MDS) plot of the Bray-Curtis dissimilarity matrix coloring study participants by relative abundance of *Firmicutes*, indicating an absence of marked stratification. Similar homogeneous distributions of other dissimilarity metrics on the MDS plot are available in Additional File 1: Figure S2.

### Associations of non-genetic variables with gut microbiome parameters

Demographic, lifestyle and environmental variables were collected via a detailed questionnaire, while biochemical parameters were measured in blood samples. Correlations between dietary consumption parameters and gut microbiome have previously been investigated in the MI cohort [29]. We considered an additional 274 variables and filtered them based on prevalence, missingness and collinearity, resulting in a final number of 110 variables to be included in association analyses (see Methods). Figure 1 outlines the six categories of non-genetic variables considered and shows representative examples. The full list with a detailed description of the tested variables is provided in Additional File 2: Table S1.

To investigate the potential impact of relevant demographic, social, behavioral, nutritional and medical data on the fecal microbiome, we searched for associations of diversity metrics and individual taxa with the 110 non-genetic variables selected above using Spearman rank testing (Additional File 2: Table S2). In total, 25 variables were significant, with on average 15 of them associated with each α-diversity metric (Additional File 1: Figure S3A) in univariate tests. Five variables (age, level of ALT, glomerular filtration rate, having breakfast and eating in fast-food restaurants) were significant (FDR < 0.05) for all α-diversity metrics (Additional File 1: Figure S3B). We then used ANOVAs to test these in multivariable models, also including four dietary variables: consumption of raw fruits, fish, fatty sweet products and sodas (which were previously found to be significantly associated with α-diversity in the same study population [29]). Only age and the levels of alanine aminotransferase (ALT), a liver enzyme whose elevated plasma levels indicate liver damage, remained significant in these analyses (Figure 3 and Additional File 2: Table S3). Simpson’s diversity index was positively associated with age and negatively associated with ALT levels, as shown in Additional File 1: Figure S4A and Additional File 1: Figure S4B.

**Figure 3.**
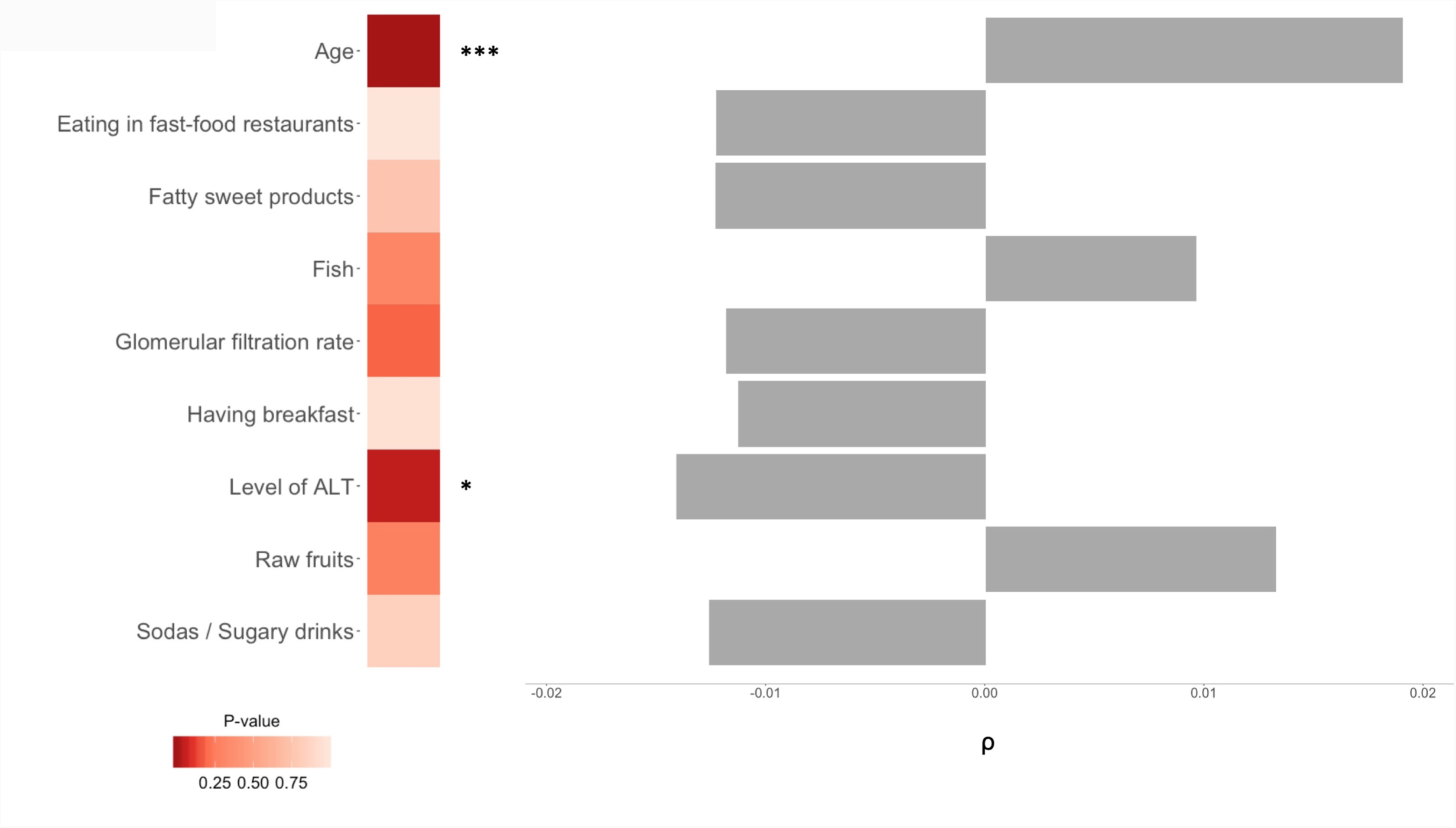
Association of non-genetic variables with α-diversity metrics. Significant variables from the univariate test and their Spearman ρ values (right hand side). Heatmap represents the ANOVA’s p-values from the multivariable test and asterisks denote statistical significance (p< 0.001: ***; p < 0.01: **; p < 0.05: *).

We then investigated the impact of non-genetic variables on the β-diversity indexes, running PERMANOVAs for the 110 variables. PERMANOVA tests a multivariate model where distance matrix is a response variable. The results of these test are presented in Additional File 2: Table S4. A total of 35 factors were significantly associated (FDR < 0.05) in univariate tests with, on average, 24 being associated with each β-diversity index (Additional File 1: Figure S5A). Fifteen factors were significant for all 3 β-diversity metrics (Additional File 1: Figure S5B). Those were then tested in multivariable models, also including raw fruit consumption (which was previously found to be significantly associated with β-diversity in our study population [29]) and reran PERMANOVAs. A total of 10 factors remained significant in the final models (Figure 4 and Additional File 2: Table S5). Of these, age, sex and plasma levels of ALT were the strongest associated factors. Also significant were chicken pox vaccination, having breakfast, having lunch, diastolic blood pressure, consumption of raw fruits, decreased or increased appetite and medical record of tooth extraction. Sex and age were able to explain the biggest portion of the observed variance of all the significantly associated variables, albeit with small individual coefficients of correlation (R^2^ < 0.01, Figure 4). We then calculated the cumulative explained variance of Bray-Curtis dissimilarity by using all the non-genetic variables available. This analysis revealed that 16.4% of the variance can be explained by non-genetic factors (Additional File 2: Table S6).

**Figure 4.**
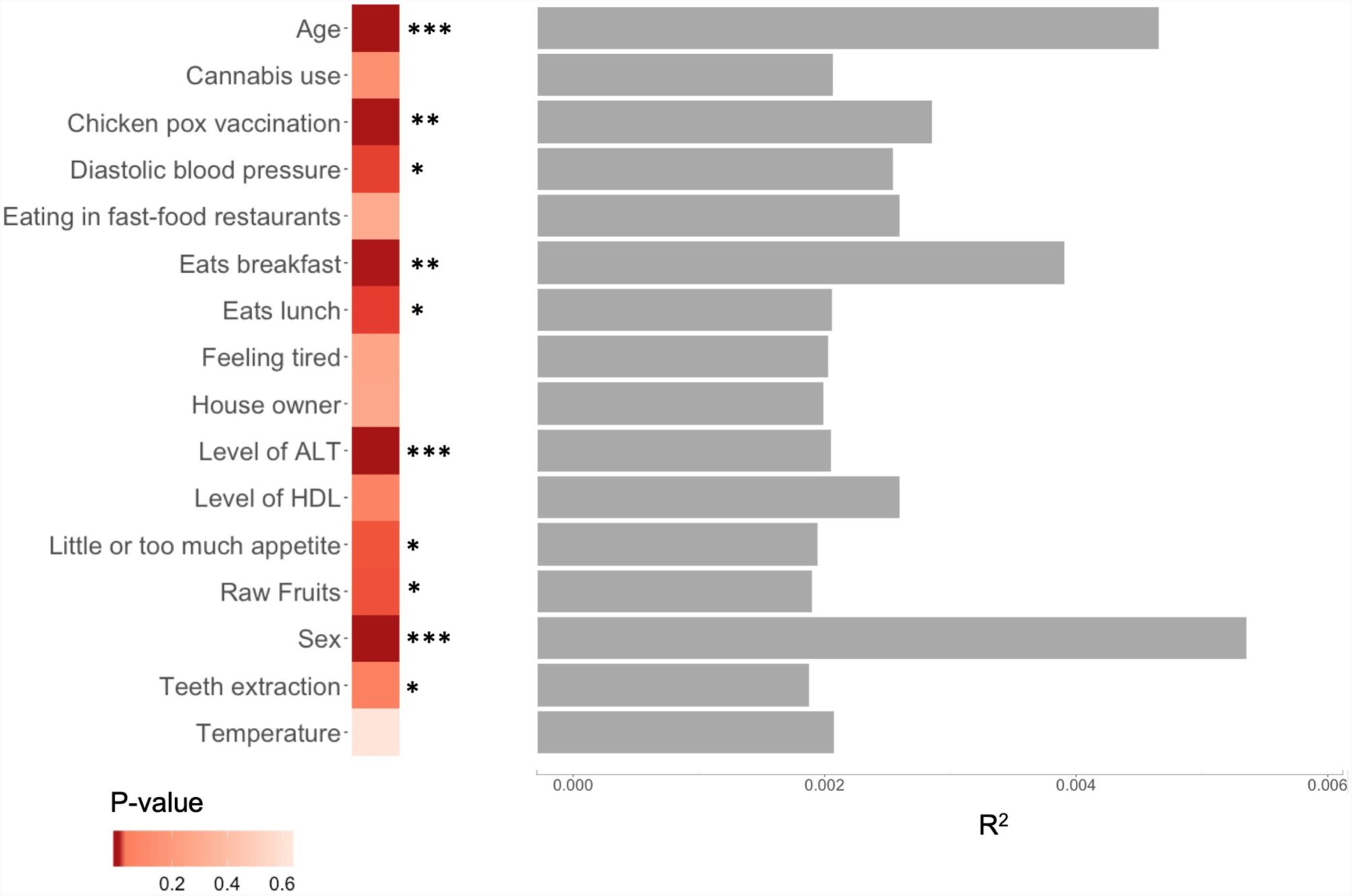
Association of non-genetic variables with β-diversity metrics. Significant variables from the univariate test and their R^2^ values (right hand side). Heatmap represents the PERMANOVA’s p-values from the multivariable test and asterisks denote statistical significance (p< 0.001: ***; p < 0.01: **; p < 0.05: *).

Next, we searched for associations between demographic and environmental variables and individual taxa. We used multivariate association with linear models to search for associations between the 110 factors discussed above and 475 taxa that were observed in more than 10% of study participants. The full list of tested taxa is available in Additional File 2: Table S7. Table 1 shows the only three significant associations (FDR corrected p-value < 0.05). We observed associations of age with the *Comamonadaceae* family and the *Schlegelella* genus, and of consumption of mineral supplements with the *Clostridium papyrosolvens* species.

**Table 1.** Significant associations of non-genetic variables with individual taxa.

| Covariate | Taxa | Prevalence | Coefficient | P-value | Q-value |
| --- | --- | --- | --- | --- | --- |
| Age | *Comamonadaceae* | 36.8% | $3.99 \times 10^{-4}$ | $3.09 \times 10^{-9}$ | $5.89 \times 10^{-5}$ |
| Age | *Schlegelella* | 29.6% | $3.32 \times 10^{-4}$ | $5.48 \times 10^{-6}$ | $3 \times 10^{-2}$ |
| Consumption of mineral supplements | *Clostridium papyrosolvens* | 13.8% | $2.44 \times 10^{-2}$ | $8.32 \times 10^{-7}$ | $4.72 \times 10^{-3}$ |

Data plots showing positive correlations of the three identified associations are presented in Additional File 1: Figure S6A-C.

### Association of human genetic variants with gut microbiome parameters

We next searched for potential associations between human genetic variants and gut microbiome diversity, using a GWAS framework. We included in the regression models all the statistically significant demographic and environmental variables identified above, for each respective phenotype. The full list of all the covariates used, including the first two principal components of the genotyping matrix, is available in Additional File 2: Table S8.

We performed GWAS using the four α-diversity metrics and the three β-diversity indexes as phenotypic outcomes. We did not observe any statistically significant association upon correction for the number of polymorphisms and of phenotypes tested (P_α-threshold_ < 1.25 × 10^−8^ and P_β-threshold_ < 1.67 × 10^−8^) (Figure 5A and Additional File 1: Figure S7; Figure 5B and Additional File 1: Figure S8). The quantile-quantile plots and lambda values, assessing the false positive rate and genomic inflation rate for all genome-wide analyses are shown in Additional File 1: Figure S9 and Figure S10. We then attempted to replicate the previously published associations between specific SNPs and β-diversity, by relaxing the genome-wide significant threshold [19, 20, 21]. Upon correction for the 66 SNPs considered (P_threshold_ < 0.05/66), none was significantly associated (Additional File 2: Table S9).

**Figure 5.**
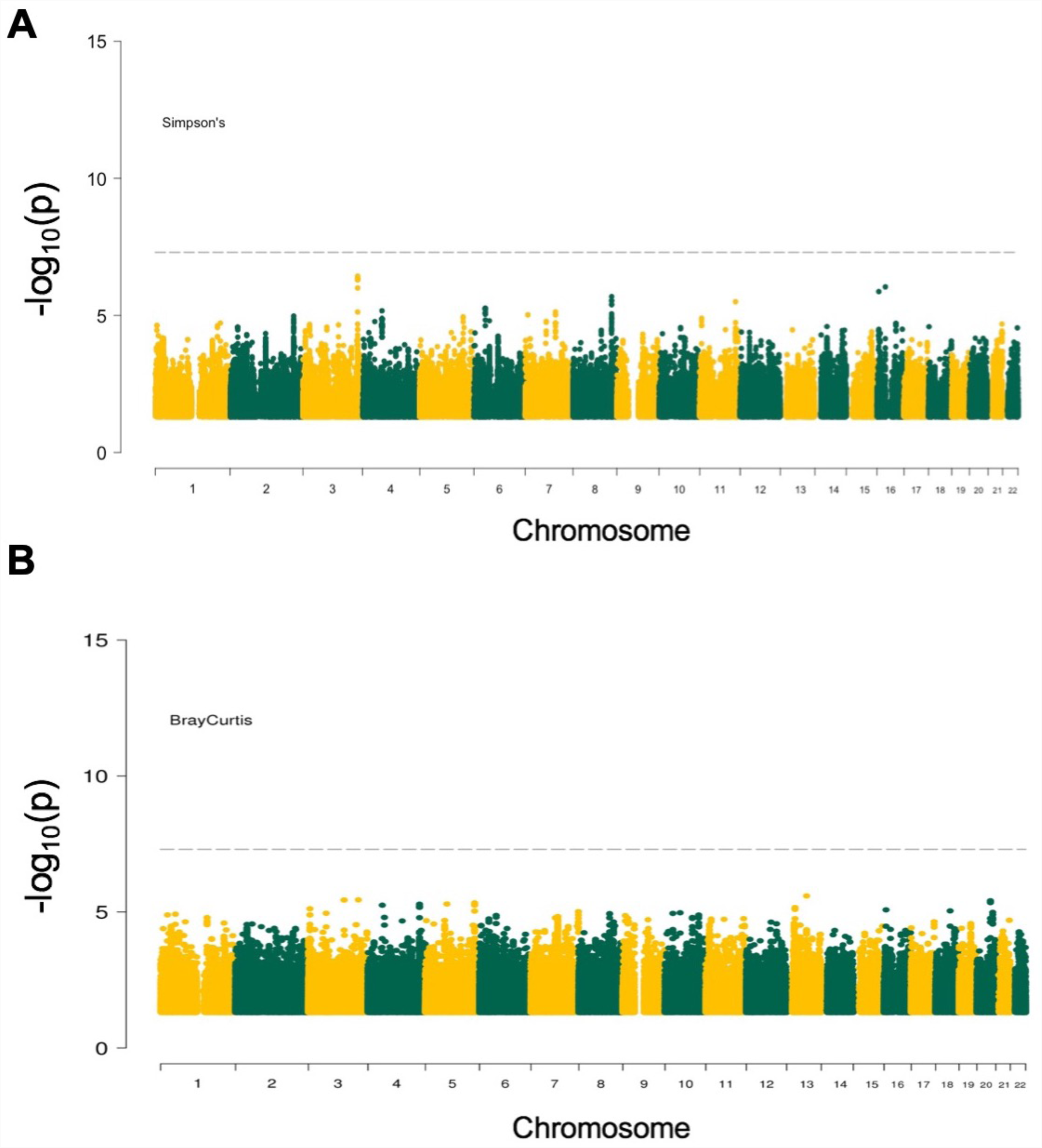
Results of genome-wide association study between host genetic variants and microbiome diversity metrics. (A) Manhattan plot for Simpson’s diversity metric (representative α-diversity metric). The dashed horizontal line denotes the genome-wide significance threshold (P_α-threshold_ < 1.25 × 10^−8^). (B) Manhattan plot for Bray-Curtis dissimilarity matrix (representative ß-diversity index). The dashed horizontal line denotes the genome-wide significance threshold (P_β-threshold_ < 1.67 × 10^−8^).

We also used a GWAS approach to search for associations between the abundance of individual taxa and human genetic variation. We used a quantitative phenotype (non-zero log-transformed relative abundance) and a binary phenotype (presence vs. absence) for each taxon. After correction for the number of polymorphisms and of phenotypes tested, we did not observe any statistically significant signal. A total of 170 suggestive associations (P_SuggestiveThreshold_ < 5 × 10^−8^) were detected with the quantitative phenotype of 53 taxa, and 65 suggestive SNPs were detected with the binary phenotype of 23 taxa. The lists of these SNPs and their association p-values are available in Additional File 2: Table S10 and Additional File 2: Table S11, respectively.

We also imputed HLA and KIR alleles and tested them for association with all the considered phenotypes, observing no significant associations (Additional file 1: Figure S11 and association summary statistics results available).

We then attempted to replicate associations for the SNPs previously reported to be associated with individual taxa (Additional File 2: Table S12) [19, 20, 21, 22, 25, 27]. Only 13 out of 336 SNPs passed the corrected nominal significance threshold (P_threshold_ < 1.49 × 10^−4^, i.e. 0.05/336) for association with a quantitative phenotype. Of these, 9 were concordant at the phylum level with the original report (i.e. the strongest associated taxon in our study belonged to the same phylum as the previously observed association). For binary phenotypes, 10 SNPs passed the corrected nominal significance threshold, including 2 that were concordant at the phylum level.

## DISCUSSION

We investigated the potential influence of demographic, environmental, clinical and genetic factors on the fecal microbiome composition in 858 unrelated healthy individuals of French descent. The *Milieu Intérieur* cohort is particularly well suited for such a comprehensive assessment [30]. The study participants have a homogeneous genetic background, live in the same region and are evenly stratified by sex and age, which provides an excellent opportunity to search for unique determinants of gut microbiome diversity.

First, we used the rich data collected through questionnaires that gathered detailed medical history as well as lifestyle and socio-demographic information. We also considered laboratory results that could indicate underlying physiological differences (e.g. levels of hemoglobin, glucose, hepatic transaminases, etc.). We searched for potential association of these variables with several α- and β-diversity metrics of the gut microbiome, as well as with quantitative and binary phenotypes derived from the detected abundance of individual microbial taxa.

As the MI cohort was designed to better understand healthy immunity, strict criteria were used during enrollment to exclude individuals with chronic medical conditions. Similarly to other studies in healthy individuals, the distribution of major phyla was in the same range as observed before (Additional file 2: Table S13). The use of prescription drugs, on the other hand, was very limited among MI participants. In fact, the final set of 110 non-genetic variables contained only one drug-related variable (“on any type of medication”). Even the use of over-the-counter drugs, such as proton pump inhibitors, was observed in less than 1% of the individuals (i.e. only in 4 individuals). The potential impact of drugs on the gut microbiome, suggested by previous studies [11, 16, 18], was therefore not evaluated in our study.

The influence of dietary variables on the gut microbiome has already been evaluated in the MI cohort [29]. Increased α-diversity was found to be associated with foods generally considered as healthy (fruits, fish), while a decrease was associated with foods for which limited consumption is generally recommended (e.g. fried products). Dissimilarity measure by β-diversity level was driven by consumption of raw fruits, fried products, ready-cooked meals, and cheese [29]. In the current analysis, we focused our attention on additional environmental influences, lifestyle variables and biochemical measurements. Age showed a strong positive association with α-diversity in all models, whereas sex and BMI did not show any consistent association. Interestingly, we replicated a correlation between higher plasma levels of alanine aminotransferase and lower microbiome diversity (previously also observed in a Belgian cohort, but not replicated in a Dutch study population [16]). The causality of the observed correlation is unclear. Indeed, much work is still needed to get a better understanding of the interplay between the gut microbiome and liver disease [31].

In the analysis of β-diversity indexes, we identified ten factors that were significant in the multivariable PERMANOVA models. In line with previous reports [6, 14, 26], we observed sex and age as the strongest influencers on all β-diversity indexes, with the lowest association p-values and highest proportion of variance explained by these factors. As other co-variates, such as environmental and host-extrinsic, are also known to impact the overall composition [32], we identified factors related to medical history (in particular chicken pox vaccination and teeth extraction), blood measurements (ALT levels and diastolic blood pressure) and lifestyle (such as tendency to have breakfast or lunch and variable appetite) having mild, yet significant, correlations with β-diversity in MI cohort. We also confirmed the independent effects of diet, in particular the consumption of raw fruits [29]. Interestingly, we could not confirm any significant association between BMI and microbiome diversity, in contrast to the recent population-based observations in the FGFP study [16]. This apparent contradiction could be partly explained by the MI study design [30]: the careful selection of healthy individuals resulted in a more limited distribution of BMI values among study participants (mean ± SD: 24.26 ± 3.26 kg/m^2^; min 18.59 and max 32). This ascertainment bias reduced our power to detect potential correlations between more extreme BMI values and microbiome diversity measurements [33]. Furthermore, an estimation of the explained variance in β-diversity metrics demonstrated a small individual effect of each variable (Additional File 2: Table S4), which together explained 16.4% of the variance. This is concordant with previous reports, where a similar proportion of variance (18.7% [16], 16.4% [17| and 20% [19]) could be explained by demographic and environmental factors. In contrast to what we observed in the MI cohort, prescription medication explained an important fraction of the variance in these other studies (up to 10% [17]), attesting to the uniqueness of our healthy study sample.

In our exploration of variables potentially associated with individual taxa, we observed a strong positive correlation between age and the *Schlegelella* genus (as well as the family it belongs to: *Comamonadaceae*). This family is very diverse, and its members have been observed both in man-made environments (various clean or polluted soils and waters) and in animals or human clinical samples [34]. The epidemiological or clinical relevance of this newly observed association is unknown. We also found an association between *Clostridium papyrosolvens*, belonging to the *Clostridia* class and *Firmicutes* phylum, and the oral intake of mineral supplements. *Clostridium papyrosolvens* is an anaerobic bacterium that is involved in the degradation of diverse carbohydrates (such as cellulose, arabinose and glucose) [35] and could thus play a role in modulating the individual glycemic response.

Our in-depth investigation of demographic, environmental and clinical variables allowed us to identify factors that are associated with various measures of gut microbiome composition. Including them as covariates in genome-wide association studies increased our power to potentially detect true genetic effects. However, after correction for multiple testing, we did not observe any statistically significant associations. This was the case for a total of 7 different α- and β-diversity metrics and for 475 individual taxa, tested either as quantitative or as binary phenotypes. We also attempted to replicate the previously reported associations between human polymorphisms and gut microbiome composition at the β-diversity or the taxonomic levels [19, 20, 21, 22, 25, 27]. None of the variants associated with β-diversity metrics replicated. For individual taxa, replication at the phylum level was successful for 2 SNPs using binary phenotypes (presence vs. absence of the phylum) and for 9 SNPs using quantitative phenotypes (abundance). Of these, only one signal was replicated at the family level: the association between rs7856187 and *Lachnospiraceae* [27]. Of note, the only SNP that was significant in a recent meta-analysis [20], rs4988235, did not show any association in our study (Additional File 2: Table S12).

## CONCLUSIONS

Our study provides an in-depth investigation of potential demographic, environmental, clinical and genetic influences on the diversity of the fecal microbiome in healthy individuals. We identified variables associated with overall microbiome composition and with a small number of individual taxa, explaining a non-negligible fraction of microbiome diversity in healthy individuals in the absence of drug treatment. The lack of any significant results in the genome-wide association analyses, on the other hand, indicates that common human genetic variants of large effects do not play a major role in shaping the gut microbiome diversity observed in healthy populations. Future studies should include larger sample sizes and a more comprehensive evaluation of human genetic variation, including rare and structural variants not captured by genotyping arrays. Evaluation of the environmental effects should be optimized for example by longitudinal tracking of study participants. Lastly, large-scale microbiome and genomic data should be pooled across cohorts, as recently proposed [36], to accelerate discovery in the field of human-microbiome interactions.

## METHODS

### The *Milieu Intérieur* cohort

The 1,000 healthy donors of the *Milieu Intérieur* cohort were recruited by BioTrial (Rennes, France). The cohort is stratified by sex (500 men, 500 women) and age (200 individuals from each decade of life, between 20 and 70 years of age). Participants were selected based on stringent inclusion and exclusion criteria, detailed elsewhere [30]. Briefly, they had no evidence of any severe/chronic/recurrent medical conditions. The main exclusion criteria were seropositivity for human immunodeficiency virus or hepatitis C virus; travel to (sub-) tropical countries within the previous 6 months; recent vaccine administration; and alcohol abuse. Subjects were excluded if they were on treatment at the time or were treated in the three months preceding enrolment with, nasal, intestinal or respiratory antibiotics or antiseptics. Volunteers following a specific diet prescribed by a doctor or dietician for medical reasons (calorie-controlled diet or diet favouring weight loss in very overweight patients, diets to decrease cholesterol levels) and volunteers with food intolerance or allergy were also excluded. To avoid the influence of hormonal fluctuations in women during the peri-menopausal phase, only pre- or post-menopausal women were included. To minimize the influence of population substructure on genomic analyses, the study was restricted to individuals of self-reported Metropolitan French origin for three generations (i.e., with parents and grandparents born in continental France). Fasting whole blood samples were collected from the 1,000 participants in lithium heparin tubes between September 2012 and August 2013.

### Fecal DNA extraction and amplicon sequencing

Human stool samples were produced at home no more than 24 hours before the scheduled medical visit and collected in a double-lined sealable bag with the outer bag containing a GENbag Anaer atmosphere generator (Aerocult, Biomerieux), used to maintain anaerobic conditions, and an anaerobic indicator strip (Anaerotest, Merck Millipore) to record the strict maintenance of the anaerobic atmosphere. Upon reception at the clinical site, the fresh stool samples were aliquoted and stored immediately at −80°C. DNA was extracted from stool as previously published [37, 38]. DNA quantity was measured with Qubit using broad range assay. Barcoding polymerase chain reaction (PCR) was carried out using indexed primers targeting the V3-V5 region of the 16S rRNA gene as described in [39]. AccuPrime™ Pfx SuperMix (Invitrogen - 12344-040) was used to perform the PCR. PCR mix was made up of 18 µL of AccuPrime™ Pfx SuperMix, 0.5 µL of both V3-340F and V5-926R primers (0.2 µM) and 1 µL of DNA (10 ng). PCR was carried out as follow: 95°C for 2 min, 30 cycles of 95°C for 20 sec, 55°C for 15 sec, 72°C for 5 min and a final step at 72°C for 10 min. Amplicon concentration was then normalized to 25 ng per PCR reaction using SequalPrep™ Normalization Plate Kit, 96-well (Thermo Fisher Scientific). Equal volumes of normalized PCR reaction were pooled and thoroughly mixed. The amplicon libraries were sequenced at the Institut Curie NGS platform on Illumina MiSeq using the 2*300 base pair V3 kit to 5,064 to 240,472 sequencing reads per sample (mean ± SD: 21,363 ± 19,087 reads).

### 16s sequencing data processing and identification of microbial taxa

Raw reads were trimmed using sickle [40], then error corrected using SPAdes [41] and merged using PEAR [42]. Reads were clustered into operational taxonomy units (OTUs) at 97% of identity using vsearch pipeline [43]. Chimeric OTUs were identified using UCHIME [44] and discarded from downstream analysis. Microbiome profiles obtained were normalized for sequencing depth (sequencing counts were divided to their sample size and then multiplied by the size of the smaller sample) [45]. We further checked the presence of the sequencing batch effect and principal coordinates analysis (PCoA) plot obtained at the genus level presented in the Additional File 1: Figure S12 shows random distribution of samples obtained from different sequencing batches.

Taxonomy of representative OTU sequences was determined using RDP classifier [46]. OTU sequences were aligned using ssu-align [47]. The phylogenetic tree was inferred from the OTUs multiple alignments using Fastree2 [48]. We further checked the specific taxonomic assignations identified in our study. *Schlegelella* genus was made of 15 OTUs that had a similarity score ranging from 60% to 80% with a phylogenetically close previously identified environmental bacteria *Schlegelella thermodepolymerans*. Furthermore, taxonomic assignation of *Clostridium papyrosolvens* was at obtained with 73% of accuracy.

For 138 individuals, the gut microbiome composition could not be established because of technical issues in the extraction and the sequencing steps (i.e. due to low DNA extraction yield, absence of PCR amplicons, low read counts). These were excluded from further analysis.

### Gut microbiome diversity estimates

Based on OTUs, we calculated two types of microbial diversity indicators: α- and β-diversity indexes. As estimates of α-diversity, we used Simpson’s diversity index, observed richness, Chao1 richness estimate and ACE (Abundance-based Coverage Estimator). We applied Yeo-Johnson transformation with R package VGAM [49] to normalize these phenotypes. The histograms of raw and transformed distributions are shown in Additional File 1: Figure S1A and Additional File 1: Figure S1B, respectively. As estimates of β-diversity, we used Bray-Curtis (weighed), compositional Jaccard (unweighted) and Unifrac (weighted) dissimilarity matrices. All diversity indicators were generated on non-rarefied data using the R package vegan [50], that was corrected for sequencing depth prior to indexes computation [45].

### Demographic, environmental and clinical variables

A large number of demographical, environmental and clinical variables are available in the *Milieu Intérieur* cohort [30]. These notably include infection and vaccination history, childhood diseases, health- and diet-related habits, socio-demographical variables, and laboratory measurements. After manual curation, we considered 274 variables as potentially interesting for our analyses. Of those, we removed 130 that: (i) were only variable in less than 5% of participants; or (ii) were missing in more than 10% of participants. We tested for collinearity among the remaining 144 variables using Spearman rank correlation. All pairwise correlations with a Spearman’s ρ > 0.6 or < −0.6 and a false discovery rate (FDR) < 5% were considered colinear; one variable from each pair was removed from further analysis, resulting in a final set of 110 variables (described in Additional File 2: Table S1). Of these, 39 had some missing values (<1% in 25, 1-5% in 10, 5-10% in 4 individuals), which were imputed using random forest method in the R package mice [51]. We evaluated the effects of various clinical measurements within their normal healthy range, such as those of BMI (mean ± SD: 24.26 ± 3.26 kg/m^2^) and C-reactive protein (CRP; mean ± SD: 1.99 ± 2.58 mg/L). Several symptoms of depression, such as lack of interest in doing things and poor self-image, and potentially relevant personal and family medical history information (such as route of birth delivery, immunization history with several vaccines and familial occurrence of diabetes or myocardial infarction) were investigated. Furthermore, smoking status and nutritional tendencies (such as the salt consumption habits) were kept in our analyses.

### Testing of demographic, environmental and clinical variables

We searched for associations between the 110 demographic, environmental and clinical variables selected above and the various gut microbiome phenotypes. For α-diversity indexes (Simpson’s index, observed richness, Chao1 richness estimate and ACE), we used non-parametric Spearman correlations. For β-diversity dissimilarities (Jaccard, Bray-Curtis and Unifrac matrices), we used permutational analysis of variance (PERMANOVA) with 1000 permutations. PERMANOVAs identify variables that are significantly associated with β-diversity and measure the fraction of variance explained by the factors tested. The variables that were significantly associated (Benjamini–Hochberg FDR < 0.05) with the diversity estimates in the univariable models were included in the respective multivariable models: we used multivariable ANOVAs for α-diversity and PERMANOVAs for β-diversity. We used a backward selection, i.e. we eliminated the variables that were not significant in the first multivariable model, and reran the tests iteratively until all included predictors were significant. Spearman correlations, ANOVA and PERMANOVAs tests were performed in R v3.5.1. Finally, to search for associations with individual taxa, we implemented multivariate association with linear models by using MaAsLin [52] with default parameters.

### Human DNA genotyping

As previously described [53], blood was collected in 5mL sodium EDTA tubes and kept at room temperature (18–25°) until processing. After extraction, DNA was genotyped at 719,665 single nucleotide polymorphisms (SNPs) using the HumanOmniExpress-24 BeadChip (Illumina). The SNP call rate was > 97% in all donors. To increase coverage of rare and potentially functional variation, 966 of the 1,000 donors were also genotyped at 245,766 exonic variants using the HumanExome-12 BeadChip. The variant call rate was < 97% in 11 donors, which were thus removed from this dataset. We filtered out from both datasets genetic variants based on a set of criteria detailed in [54]. These quality-control filters yielded a total of 661,332 and 87,960 variants for the HumanOmniExpress and HumanExome BeadChips, respectively. Average concordance rate for the 16,753 SNPs shared between the two genotyping platforms was 99.99%, and individual concordance rates ranged from 99.8% to 100%.

### Genetic relatedness and structure

Relatedness was detected using KING [55]. Six pairs of related participants (parent-child, first and second-degree siblings) were identified. Of those, four pairs had both genotyping and microbiome datasets and one individual from each pair, randomly selected, was removed from the genetic analyses, leaving in total 858 individuals with both genotyping and 16s rRNA gene sequencing data. The genetic structure of the study population was estimated using principal component analysis (PCA), implemented in EIGENSTRAT (v6.1.3) [56]. The PCA plot of the study population is shown in Additional File 1: Figure S13.

### Genotype imputation

As described previously [54], we used Positional Burrows-Wheeler Transform for genotype imputation, starting with the 661,332 quality-controlled SNPs genotyped on the HumanOmniExpress array. Phasing was performed using EAGLE2 (v2.0.5) [57]. As reference panel, we used the haplotypes from the Haplotype Reference Consortium (release 1.1) [58]. After removing SNPs that had an imputation info score < 0.8, we obtained 22,235,661 variants. We then merged the imputed dataset with 87,960 variants directly genotyped on the HumanExome BeadChips array and removed variants that were monomorphic or diverged significantly from Hardy-Weinberg equilibrium (P < 10^−7^). We obtained a total of 12,058,650 genetic variants to be used in association analyses.

We used SNP2HLA (v1.03) [59] to impute 104 4-digit human leukocyte antigen (HLA) alleles and 738 amino acid residues (at 315 variable amino acid positions of the HLA class I and II proteins) with a minor allele frequency (MAF) of >1%.

We used KIR*IMP [60] to impute killer-cell immunoglobulin-like receptor (KIR) alleles, after haplotype inference on chromosome 19 with SHAPEIT2 (v2.r790) [61]. A total of 19 KIR types were imputed: 17 loci plus two extended haplotype classifications (A vs. B and KIR haplotype). A MAF threshold of 1% was applied, leaving 16 KIR alleles for association analysis.

### Genetic association analyses

For single-variant association analyses, we only considered SNPs with a MAF higher then 5% (N=5,293,637). Unless otherwise stated, we used PLINK (v1.9) [62] for association testing. In all tests, we included the first two first principal components of the genotyping matrix as covariates to correct for residual population stratification. The demographic, environmental and clinical variables that were identified as significantly associated were also included as covariates in the respective analyses. A full list of covariates for each phenotype is available in Additional File 2: Table S8.

We used linear regression (within PLINK) and microbiomeGWAS [63] to test for SNP associations with α-diversity indexes and β-diversity dissimilarities, respectively. Linear regression was also used to search for associations with relative abundance of specific taxa. Only taxa present in at least 10% of individuals were tested (N=475), i.e. 8/11 (remaining/total) phyla, 16/24 classes, 20/43 orders, 50/103 families, 135/328 genera and 246/698 species. The list of all tested taxa is presented in Additional File 2: Table S7. We used logistic regression to test binary phenotypes (presence/absence of specific taxa). Here, we excluded taxa that were present in >90% of individuals, resulting in a total of 374 phenotypes (4 phyla, 8 classes, 15 orders, 38 families, 104 genera and 205 species). For all GWAS, we used a significance threshold corrected for the number of tests performed. For α-diversity (N=4): P_α-threshold_ < 1.25 × 10^−8^, for β-diversity (N=3): P_β-threshold_ < 1.67 × 10^−8^, for taxa abundance (N=475): P_taxa-linear_ < 1.05 × 10^−10^ and for presence or absence of taxa (N=374): P_taxa-logistic_ < 1.33 × 10^−10^.

## LIST OF ABBREVIATIONS

SNP: single nucleotide polymorphism;
MAF: minor allele frequency;
MI: *Milieu Intérieur*;
QQ: quantile-quantile;
LD: linkage disequilibrium;
PCR: polymerase chain reaction;
ANOVA: analysis of variance;
PERMANOVA: permutational analysis of variance;
FDR: false discovery rate;
OTU: operational taxonomy unit;
HIV: human immunodeficiency virus;
HCV: hepatitis C virus;
ACE: Abundance-based coverage estimator;
GWAS: genome-wide association study;
HLA: human leukocyte antigen;
KIR: killer-cell immunoglobulin-like receptors;
PCA: principal component analysis;
MDS: Multidimensional scaling;
PCoA: Principal Coordinates Analysis;
CRP: C-reactive Protein;
ALT: Alanine transaminase;
rRNA: Ribosomal ribonucleic acid.

## DECLARATIONS

### Ethics approval and consent to participate

The clinical study was approved by the Comité de Protection des Personnes - Ouest 6 on June 13th, 2012, and by the French Agence Nationale de Sécurité du Médicament on June 22nd, 2012 and has been performed in accordance with the Declaration of Helsinki. The study is sponsored by the Institut Pasteur (Pasteur ID-RCB Number: 2012-A00238-35) and was conducted as a single center study without any investigational product. The protocol is registered under ClinicalTrials.gov (study number NCT01699893). Informed consent was obtained from participants after the nature and possible consequences of the studies were explained.

### Consent for publication

Not applicable.

### Availability of data and material

Genotype data supporting the conclusions of this article are available in the European Genome-Phenome Archive under the accession code EGAS00001002460. Full summary association results will be made available for download from Zenodo.

### Authors’ contributions

Conception of the study: J.F., L.Q.-M., D.D., M.L.A, O.L.; Design of the study: P.S., F.H., A.B., J.F.; Acquisition of the data: P.S., S.M., V.P., C.H., C.A., J.B., E.P., M.T.; Analysis of the data: P.S., F.H., J.F.; Drafting the manuscript: P.S., J.F.; Revising the manuscript: P.S., F.H., S.M., V.P., A.B., C.H., C.A., J.B., E.P., M.T., O.L., M.L.A., D.D., L.Q.-M., J.F.

## Acknowledgements

We would like to thank to all the donors for their contribution to the study.

The *Milieu Intérieur* Consortium is composed of the following team leaders: Laurent Abel (Hôpital Necker, Paris, France), Andres Alcover (Institut Pasteur, Paris, France), Hugues Aschard (Institut Pasteur, Paris, France), Kalla Astrom (Lund University, Lund, Sweden), Philippe Bousso (Institut Pasteur, Paris, France), Pierre Bruhns (Institut Pasteur, Paris, France), Ana Cumano (Institut Pasteur, Paris, France), Caroline Demangel (Institut Pasteur, Paris, France), Ludovic Deriano (Institut Pasteur, Paris, France), James Di Santo (Institut Pasteur, Paris, France), Françoise Dromer (Institut Pasteur, Paris, France), Darragh Duffy (Institut Pasteur, Paris, France), Gérard Eberl (Institut Pasteur, Paris, France), Jost Enninga (Institut Pasteur, Paris, France), Jacques Fellay (EPFL, Lausanne, Switzerland) Odile Gelpi (Institut Pasteur, Paris, France), Ivo Gomperts-Boneca (Institut Pasteur, Paris, France), Milena Hasan (Institut Pasteur, Paris, France), Serge Hercberg (Université Paris 13, Paris, France), Olivier Lantz (Institut Curie, Paris, France), Claude Leclerc (Institut Pasteur, Paris, France), Hugo Mouquet (Institut Pasteur, Paris, France), Sandra Pellegrini (Institut Pasteur, Paris, France), Stanislas Pol (Hôpital Côchin, Paris, France), Antonio Rausell (INSERM UMR 1163 – Institut Imagine, Paris, France), Lars Rogge (Institut Pasteur, Paris, France), Anavaj Sakuntabhai (Institut Pasteur, Paris, France), Olivier Schwartz (Institut Pasteur, Paris, France), Benno Schwikowski (Institut Pasteur, Paris, France), Spencer Shorte (Institut Pasteur, Paris, France), Vassili Soumelis (Institut Curie, Paris, France), Frédéric Tangy (Institut Pasteur, Paris, France), Eric Tartour (Hôpital Européen George Pompidou, Paris, France), Antoine Toubert (Hôpital Saint-Louis, Paris, France), Mathilde Touvier (Université Paris 13, Paris, France), Marie-Noëlle Ungeheuer (Institut Pasteur, Paris, France), Matthew L. Albert (Roche Genentech, South San Francisco, CA, USA), Lluis Quintana-Murci (Institut Pasteur, Paris, France). Matthew L. Albert and Lluis Quintana-Murci are co-coordinators of the consortium. Additional information can be found at: http://www.milieuinterieur.fr/en.

## Additional Files

Additional File 1: (DOCX 3.4 MB)

Figure S1. Raw and transformed distributions and violin plots of α-diversity phenotypes.

Figure S2. Multidimensional scaling plots of Jaccard and Unifrac distance matrices.

Figure S3. Number and overlap of non-genetic variables associated with α-diversity phenotypes.

Figure S4. Correlations of age and ALT levels with Simpson’s diversity index.

Figure S5. Number and overlap of non-genetic variables associated with ß-diversity matrices.

Figure S6. Data plots showing identified correlations of non-genetic variables with three individual taxa.

Figure S7. Manhattan plots for α-diversity metrics: richness, Chao1 and ACE.

Figure S8. Manhattan plots for ß-diversity matrices: Jaccard and Unifrac.

Figure S9. QQ plots and lambda values of GWAS of α-diversity phenotypes.

Figure S10. QQ plots and lambda values of GWAS of β-diversity indexes.

Figure S11.Manhattan plot of HLA and KIR association results with all phenotypes. Figure S12. PCoA plot of samples obtained from different sequencing batches.

Figure S13. PCA plot of the genetic matrix data of MI donors.

Additional File 2: (XLSX 631 KB)

Table S1. Description of all the covariates used in the study.

Table S2. Spearman correlations of all the covariates with four α-diversity metrics. Table S3. Results of multivariable ANOVAs with α-diversity metrics.

Table S4. PERMANOVA results for all of the covariates with three β-diversity indexes. Table S5. Results of multivariable PERMANOVAs with β-diversity indexes.

Table S6. Explained cumulative variance of Bray-Curtis dissimilarity metric by all non-genetic covariates (dietary and 110 tested in this study).

Table S7. List of taxa tested for association with genetic variants.

Table S8. List of identified covariates that were used for each phenotype in addition to the first two principal components of the genotyping matrix.

Table S9. Replication of the SNPs previously reported to be associated with β-diversity.

Table S10. Nominal associations of SNPs with relative abundances of taxa.

Table S11. Nominal associations of SNPs with dichotomized taxa in the MI cohort.

Table S12. Replication of the SNPs previously reported to be associated with individual taxa. Table S13. Mean relative abundance of major phyla in MI and other studies.

