## Supplementary Figures for "A Comprehensive Assessment of Demographic, Environmental and Host Genetic Associations with Gut Microbiome Diversity in Healthy Individuals"

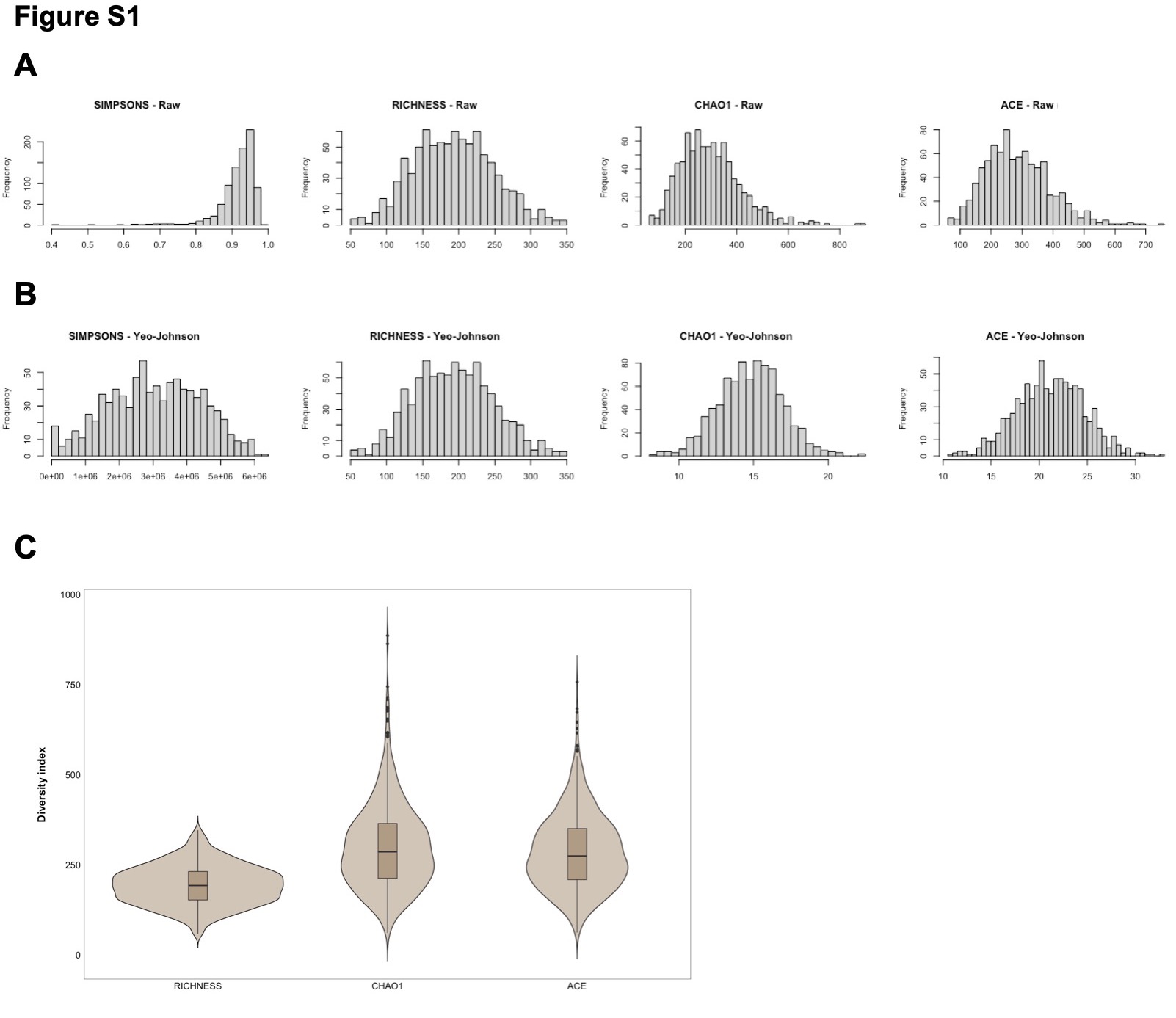

**Figure S1.** Distributions of four (A) raw and (B) Yeo-Johnson transformed α-diversity phenotypes. (C) Violin plots of distributions of α-diversity metrics – Richness, Chao1 and ACE.

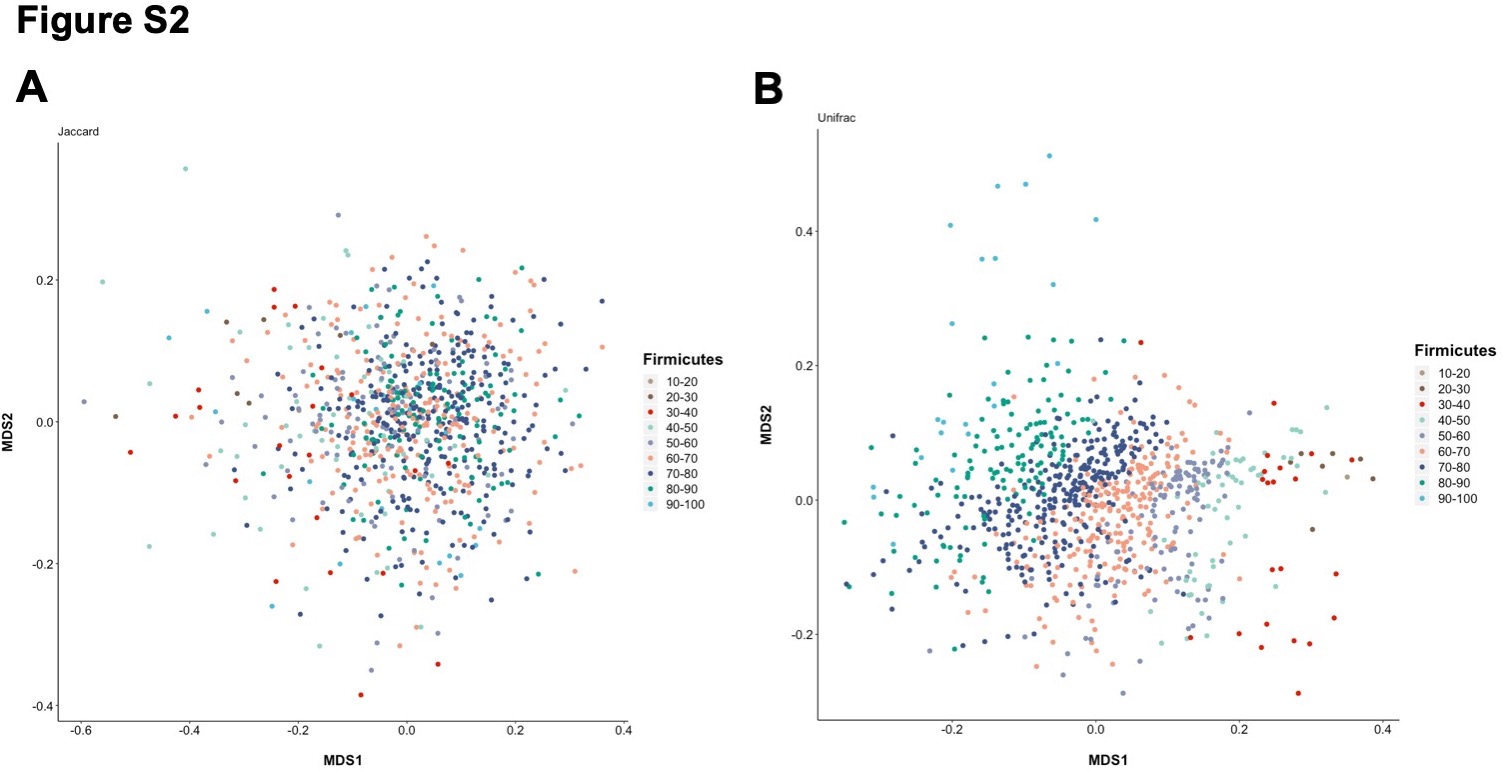

**Figure S2.** Multidimensional scaling plots of Jaccard (left hand side) and Unifrac (right hand side) distances with samples colored according to the relative abundance of *Firmicutes*.

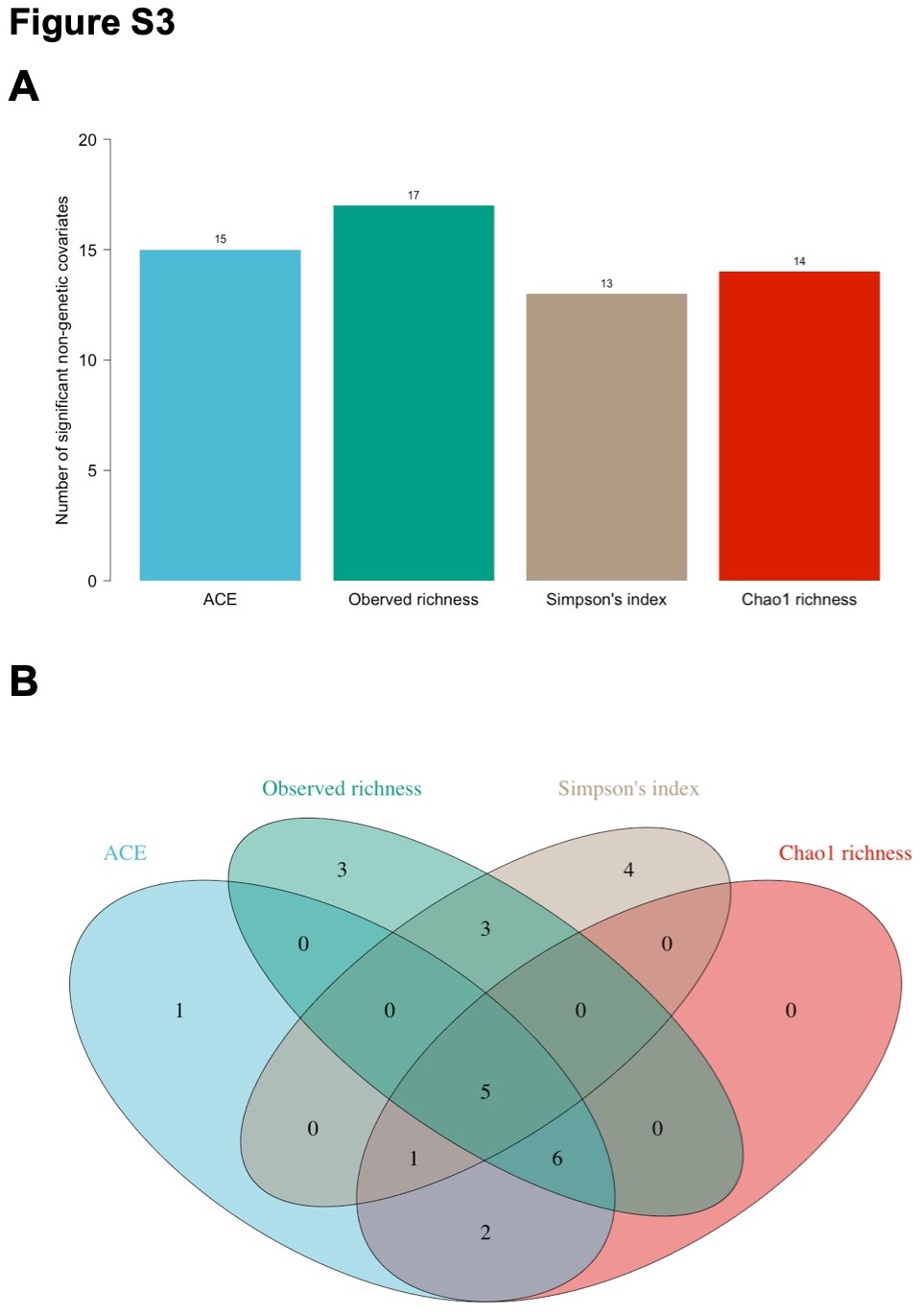

**Figure S3.** (A) Number of significant non-genetic covariates from univariate tests with four α-diversity metrics and their overlap (B).

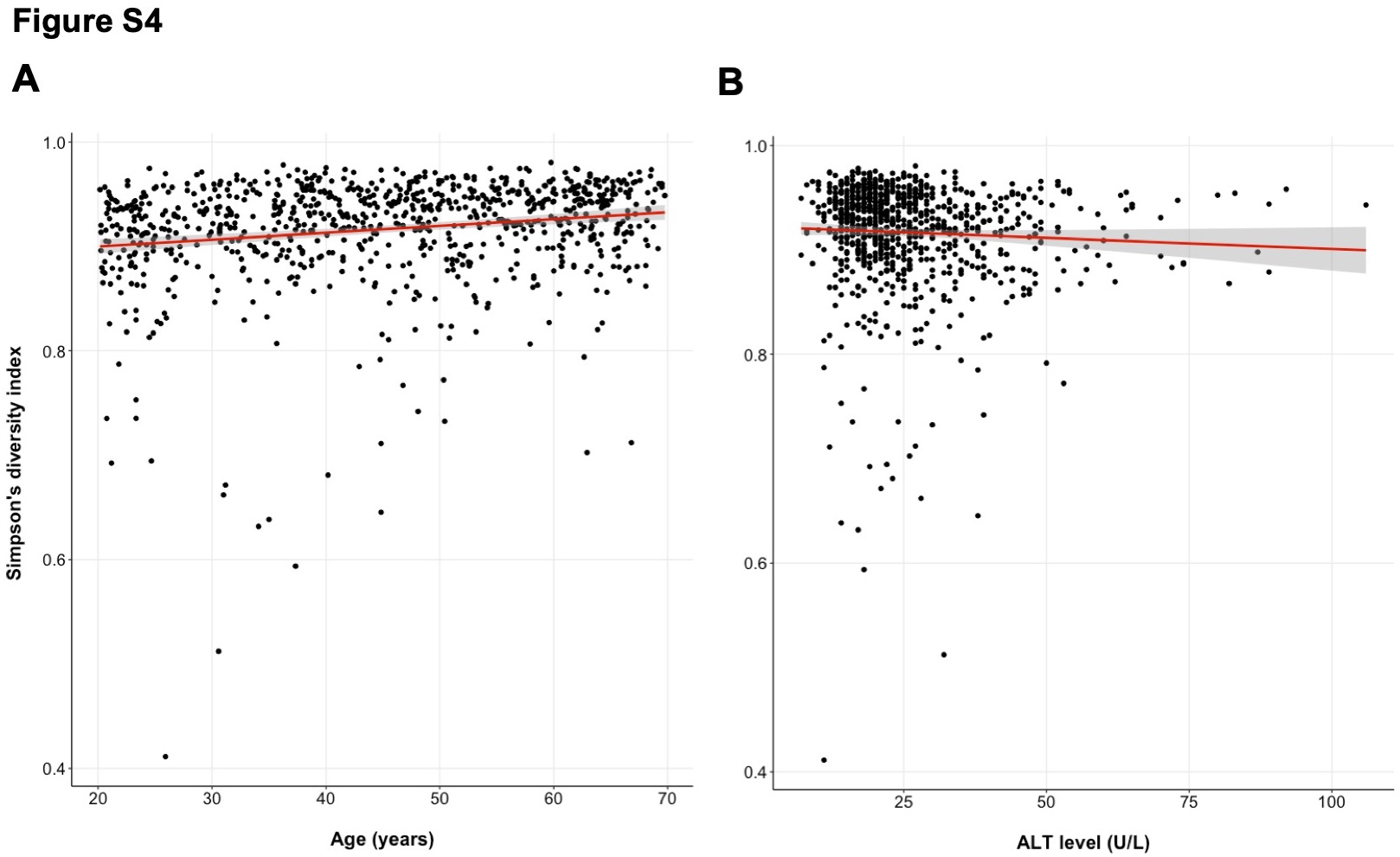

**Figure S4.** Scatter plots and the line of the best fit of Simpson’s diversity index (representative

α-diversity metric) with (A) age and (B) ALT level variables.

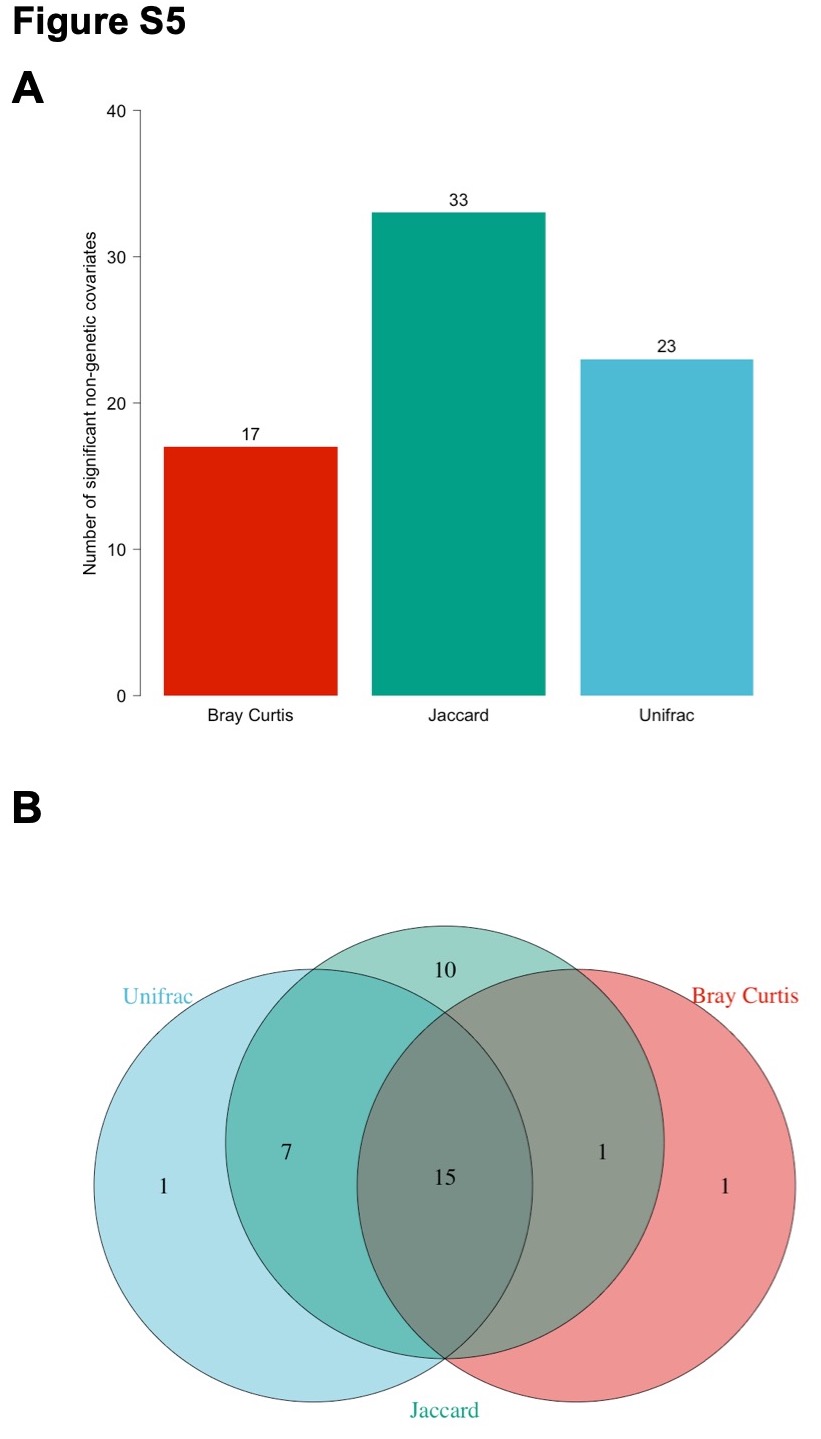

**Figure S5.** (A) Number of significant non-genetic covariates from univariate tests with three β-diversity metrics and their overlap (B).

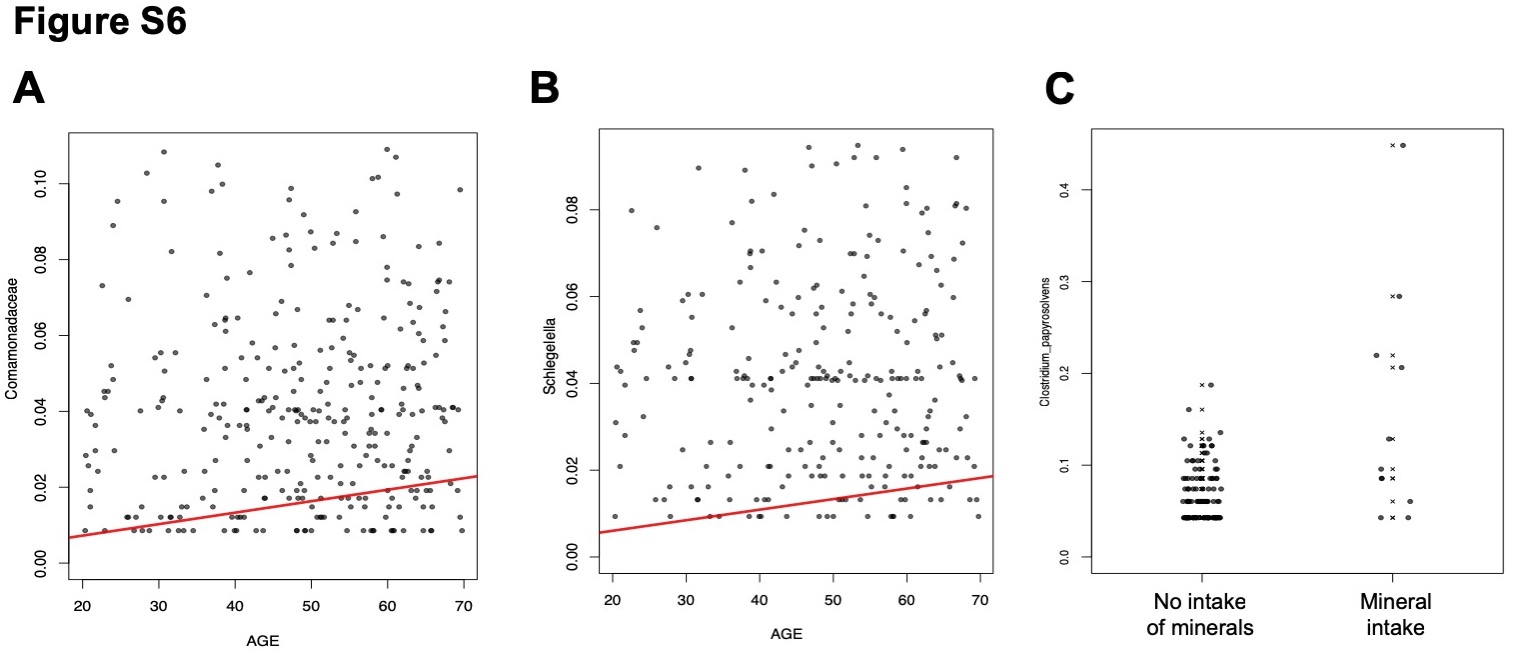

**Figure S6.** Data plots indicating association between age and (A) *Comamonadaceae*, (B)

*Schlegelella*, and the consumption of minerals with (C) *Clostridium papyrosolvens*.

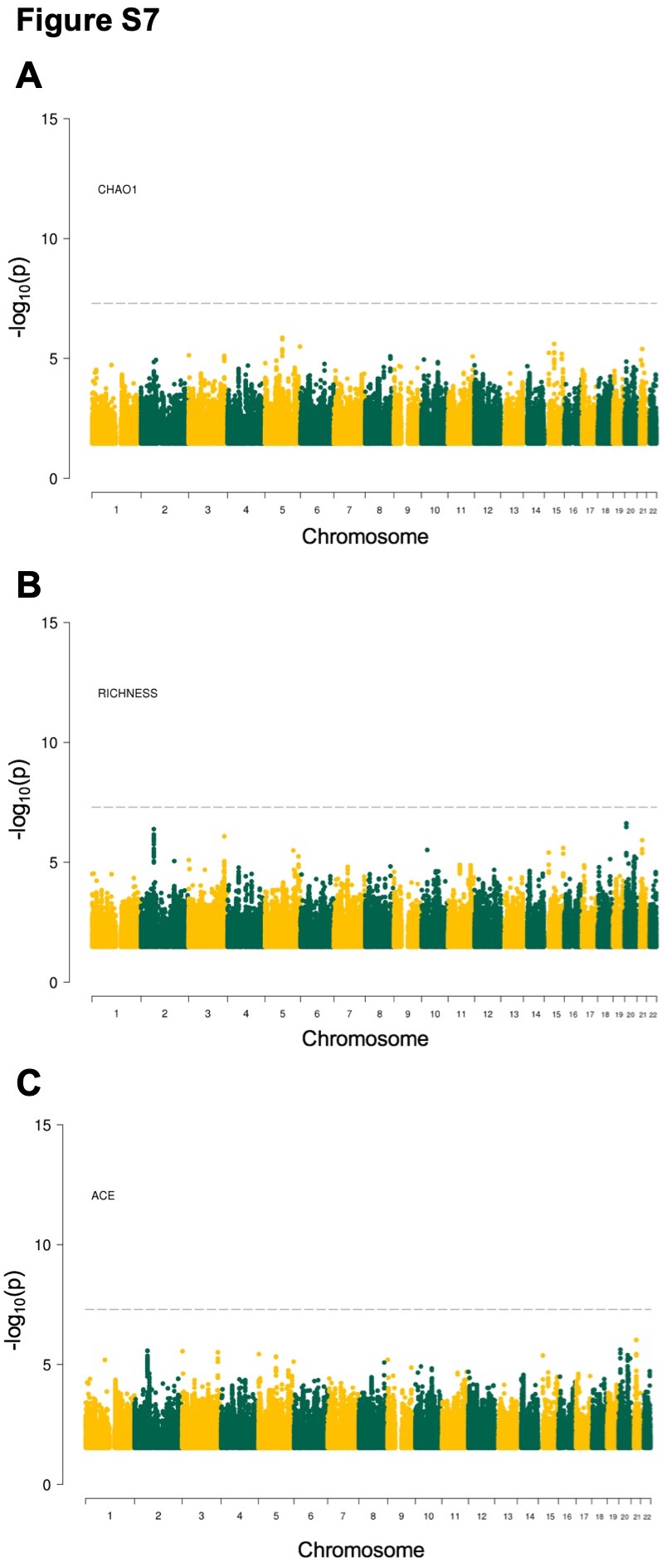

**Figure S7.** Manhattan plots showing the results of GWAS of (A) Chao1 index, (B) richness and (C) ACE. The dashed horizontal line denotes genome-wide significance threshold (P_α-threshold_ < 1.25 x 10^-8^).

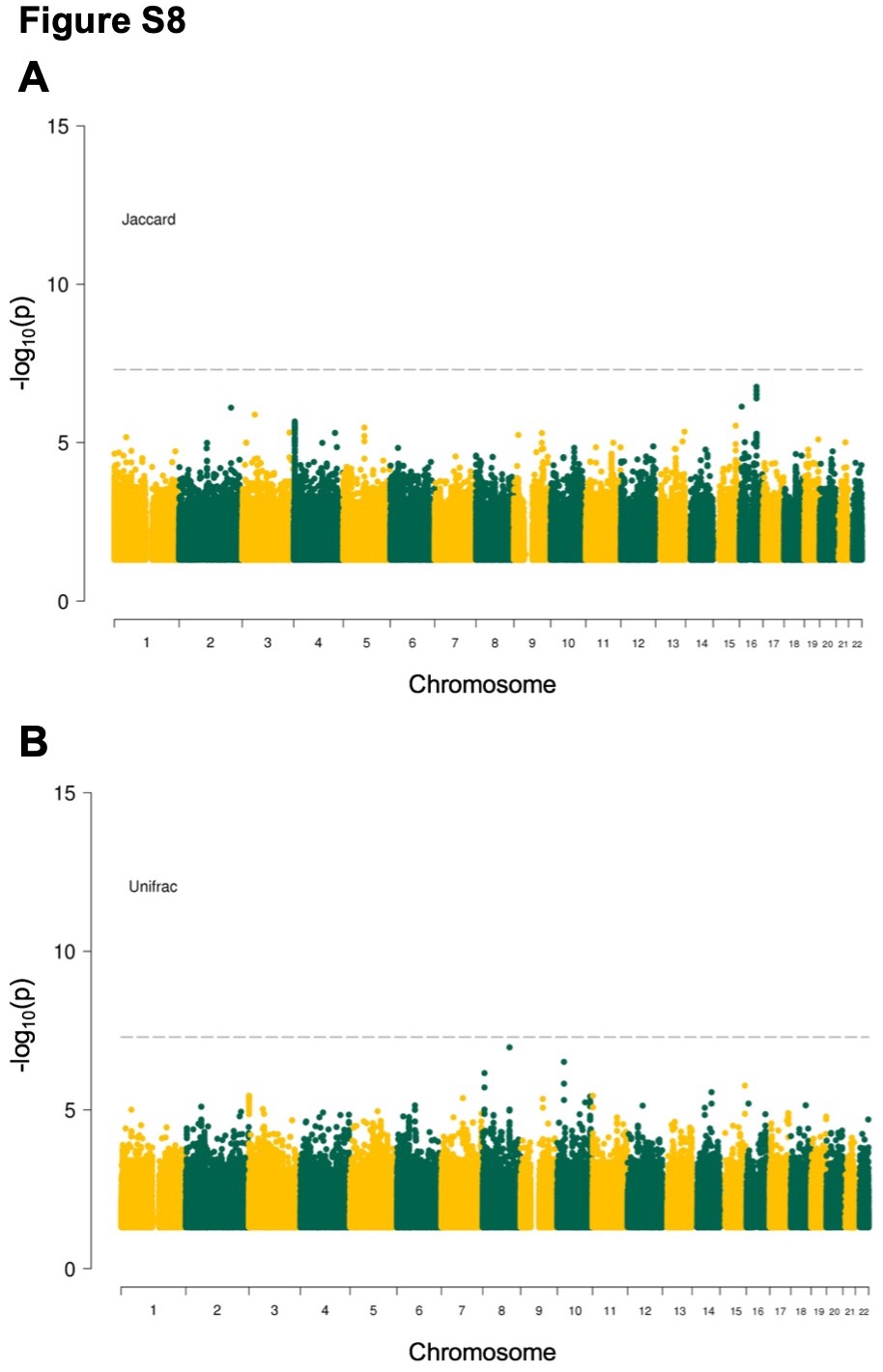

**Figure S8.** Manhattan plots showing the results of GWAS of (A) Jaccard and (B) Unifrac distances. The dashed horizontal line denotes genome-wide significance threshold (P_β-threshold_ < 1.67 x 10^-8^).

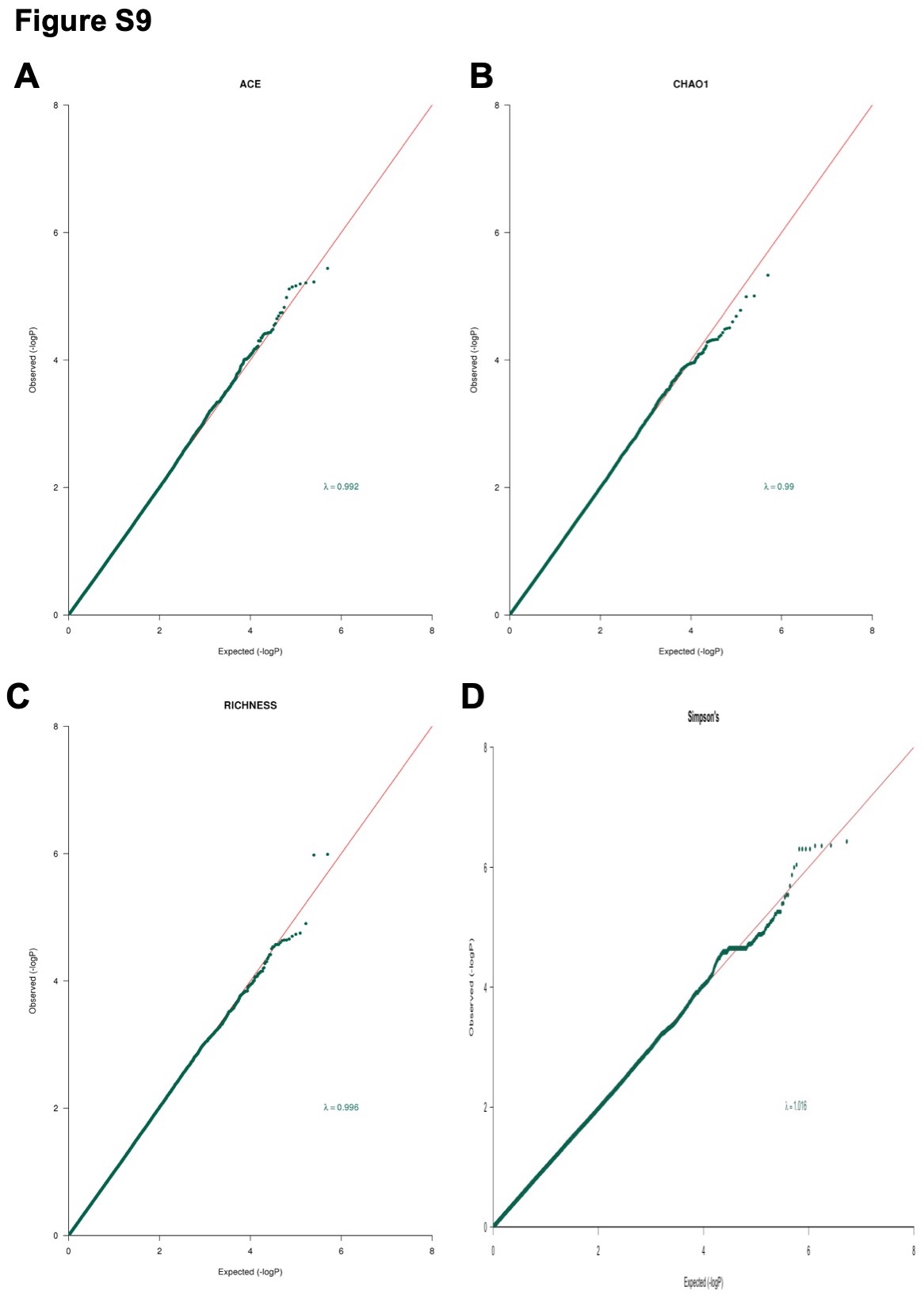

**Figure S9.** The quantile-quantile plots and lambda values of genome-wide associations preformed for (A) Simpson’s index, (B) Chao1 index, (C) richness and (D) ACE.

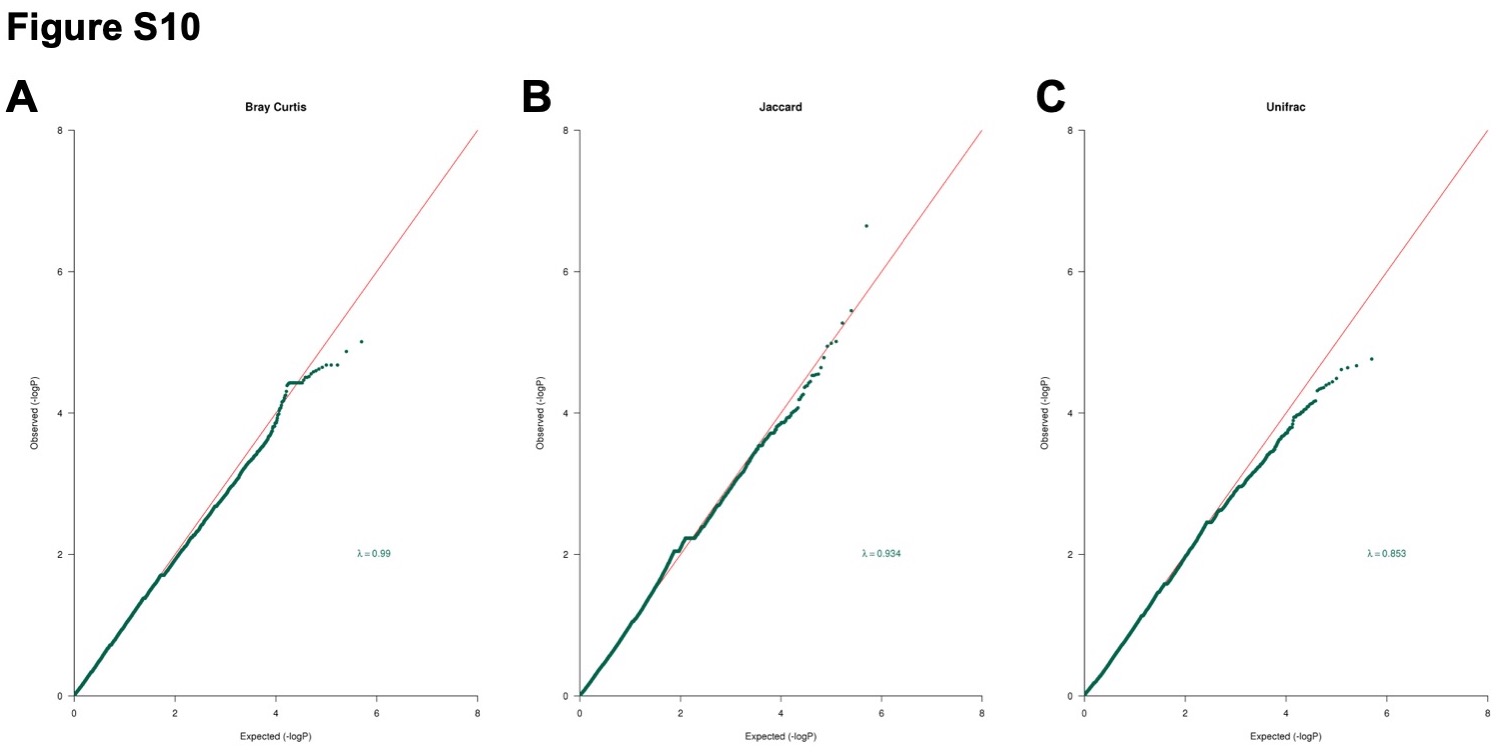

**Figure S10.** The quantile-quantile plots and lambda values of genome-wide associations preformed for (A) Bray Curtis, (B) Jaccard and (C) Unifrac distances.

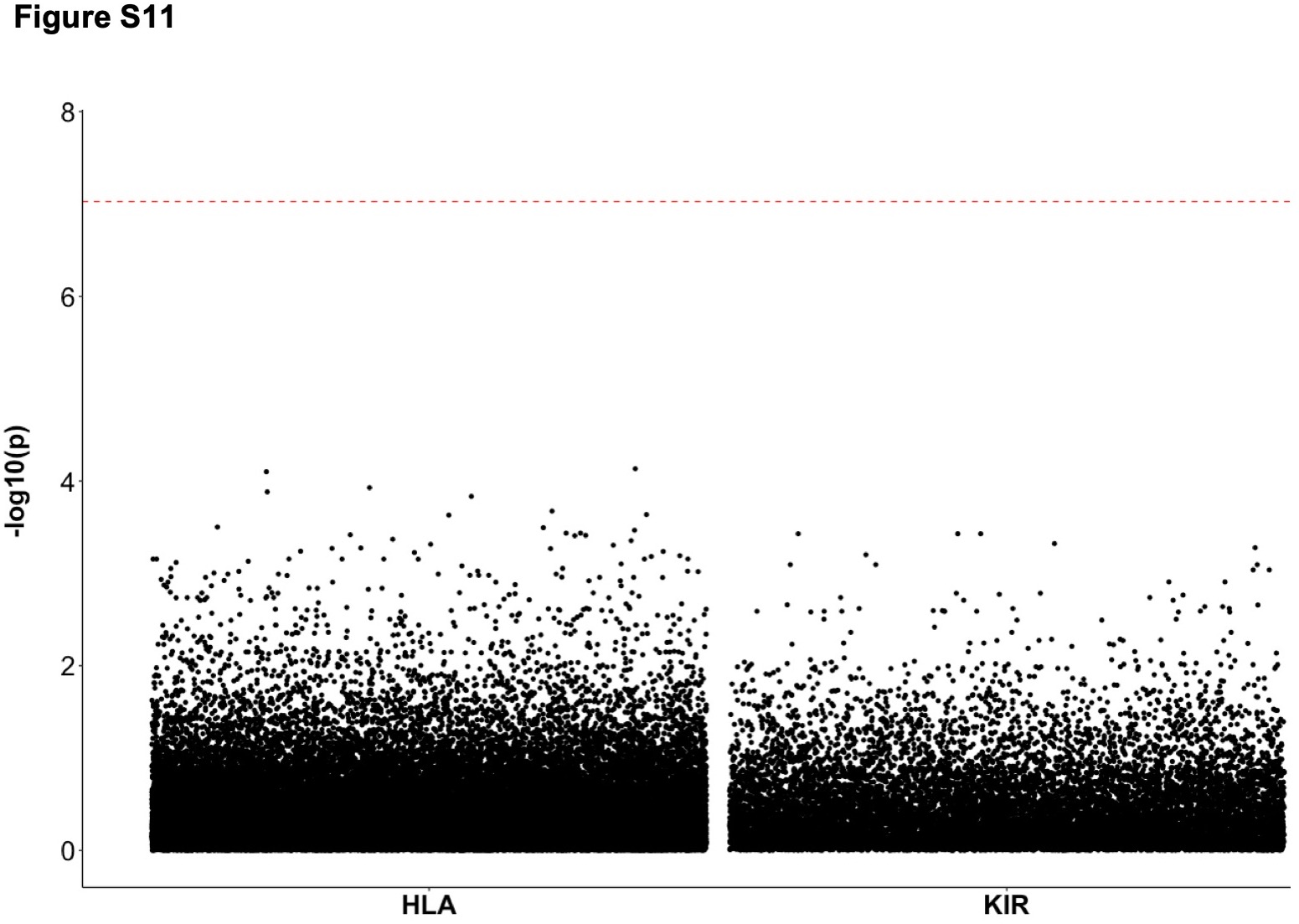

**Figure S11.** Manhattan plot of association results of all tested HLA and KIR alleles with all phenotypes (α-diversity, β-diversity, binary and quantitative taxa). The dashed horizontal line denotes significance threshold corrected for the number of tests performed (P_threshold_ < 9.42 x 10^-7^).

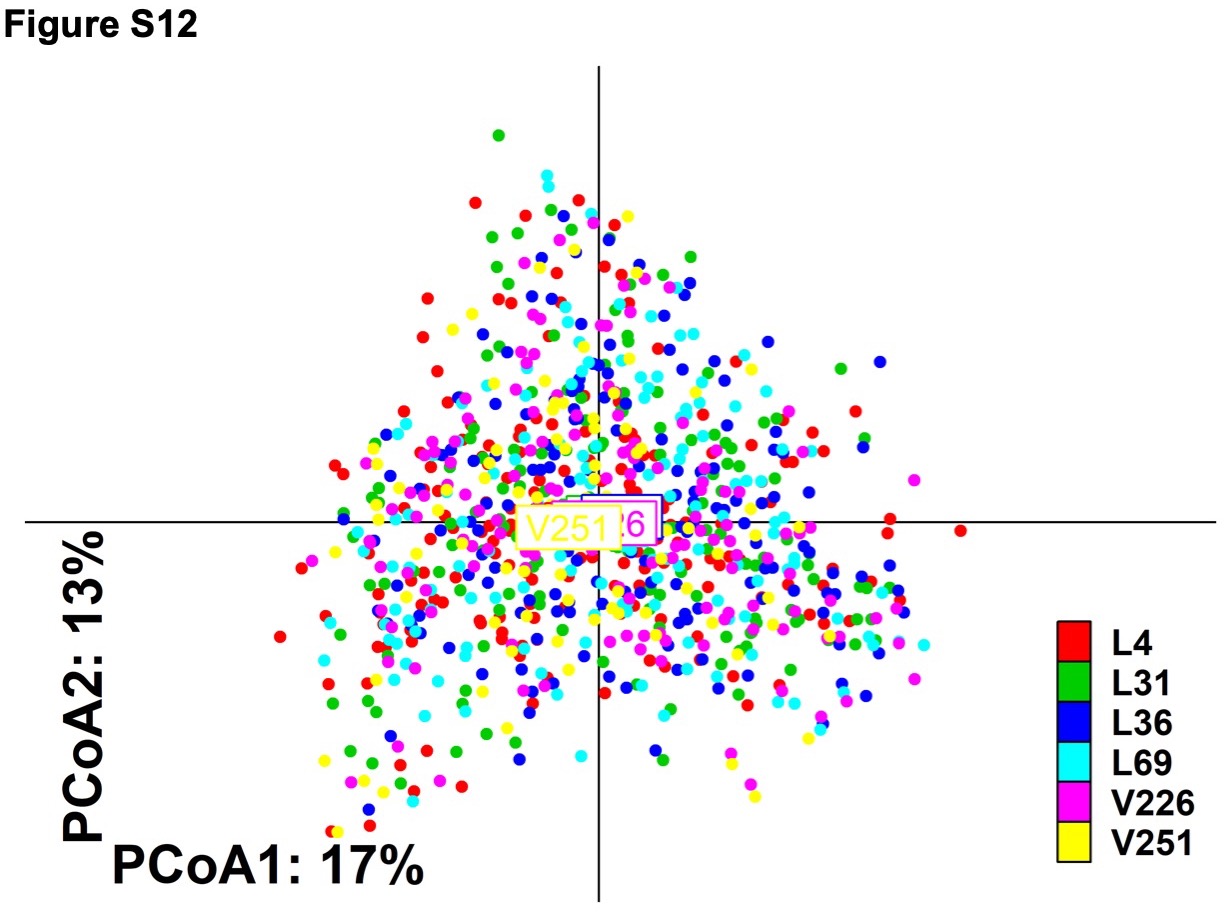

**Figure S12.** Principal Coordinate Analysis (PCoA) obtained from the genus level of MI samples deriving from different sequencing batches (indicated as L4, L31, L36, L69, V226, V251).

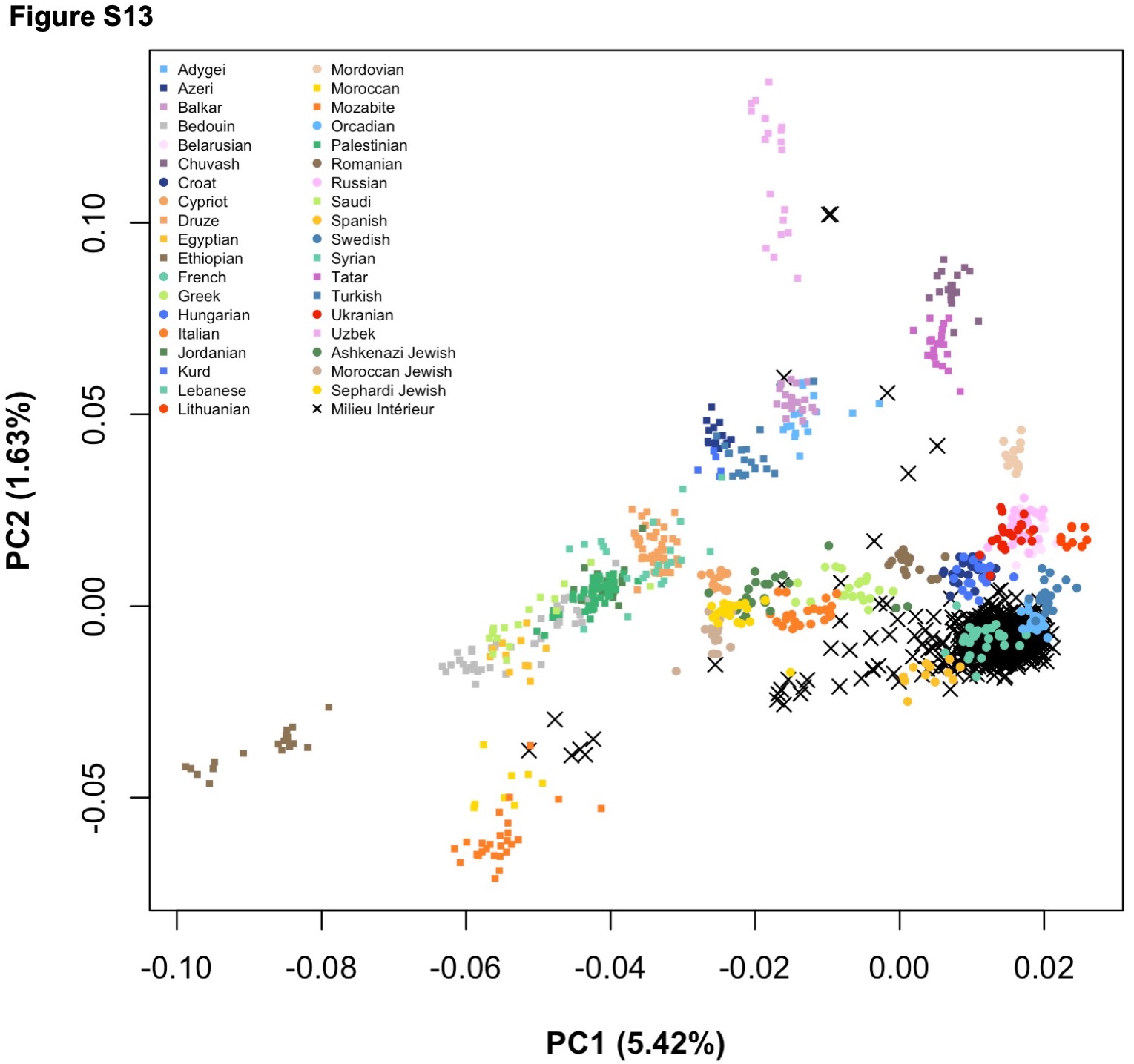

**Figure S13.** PCA plot of the genetic matrix data of MI donors. Adapted from [51].
